# The temperature dependence of binding entropy is a selective pressure in protein evolution

**DOI:** 10.1101/2025.03.23.644840

**Authors:** Rosemary Georgelin, Hannah Bott, Joe A. Kaczmarski, Rebecca Frkic, Li Lynn Tan, Nobuhiko Tokuriki, Matthew A. Spence, Colin J. Jackson

**Affiliations:** Research School of Chemistry, Australian National University, Canberra, ACT 2601, Australia; Australian Research Council Centre of Excellence for Innovations in Peptide and Protein Science, Research School of Chemistry, Australian National University, Canberra, ACT 2601, Australia; Research School of Biology, Australian National University, Canberra, ACT 2601, Australia; Michael Smith Laboratories, University of British Columbia, Vancouver, BC, V6T 1Z4, Canada; Australian Research Council Centre of Excellence in Synthetic Biology, Australian National University, Canberra, ACT 2601, Australia

**Author notes:** These authors contributed equally (Rosemary Georgelin, Hannah Bott).

## Abstract

Proteins operate through ligand and solvent interactions governed by thermodynamics, yet the enthalpy-entropy trade-offs that guide their functional evolution remain poorly understood. The LacI/GalR family (LGF) of transcription factors provides a system for examining how these trade-offs evolve over billions of years but has seldom been studied from a full thermodynamic perspective. While the evolution of ligand specificity has been well-studied, the thermodynamic determinants underlying the changes in specificity is not well understood – especially how proteins alter thermodynamic strategies to optimize affinity. By reconstructing LGF ancestors, we reveal a shift from entropy-driven binding in the most distant ancestor, to enthalpy-driven binding in the most recent ancestor and extant LacI. The most distant ancestor is characterized by the ability to bind its ligand in an open and dynamic conformation, and we propose that entropically-driven binding is driven by the presence of entropic reservoirs. This thermodynamic binding trade-off between the most distant and most recent ancestor is in accordance with the concept of ancient life that existed in a hot Earth environment, where higher temperatures enhanced entropically-driven binding. This suggests that molecular binding mechanisms evolved not just for ligand specificity, but to adapt to environmental pressures such as cooling Earth temperatures where enthalpic binding modes are favored.

## Introduction

A central goal in the study of molecular evolution is to understand the physical mechanisms that underpin phenotypic diversification as proteins evolve to adapt to challenges that are presented by a changing environment. Functional promiscuity is an important aspect of molecular evolution and is the result of non-productive activities that are under neutral selection and overlap with a protein’s native function^1–4^. Promiscuity creates a reservoir of biochemical novelty that evolution can explore under environmental selection pressures^5^, yet the thermodynamic determinants of molecular adaptation are not well understood. The binding of a protein to its molecular partners is driven by the sum of enthalpic (ΔH) and entropic (-TΔS) contributions, but it is unclear how natural selection traverses the free-energy landscape while balancing these thermodynamic parameters in molecular adaptation. An important question is whether the binding thermodynamics of chemically similar ligands are conserved over evolutionary trajectories, or do different strategies evolve to optimize enthalpic or entropic contributions to ligand binding?

The LacI/GalR family (LGF) of transcription factors (TFs) is an important system in the study of molecular evolution and biophysics. Members of the LGF are ubiquitous across bacteria and regulate the expression of metabolic genes^6^. Ligand binding in the ligand binding domain (LBD) alters the DNA-binding affinity of an N-terminal DNA-binding domain via long range allosteric interactions (DBD), coupling gene expression to nutrient availability^7,8^. The LGF regulatory domain shares homology with solute-binding proteins (SBP), which are an ancient superfamily that have been used as model systems previously to study the emergence of catalysis^9,10^, and binding specificity ^4^. The progenitor of the LGF likely emerged in the last bacterial common ancestor following the fusion of an SBP with a DNA-binding domain^11^. Owing to their ability to couple ligand concentration with organismal level phenotype via control of transcription, proteins of this family have become major protein engineering targets because of their potential use in whole cell biosensing and engineering gene regulation^12–18^.

Under the evolutionary hypothesis of fusion between an ancient SBP and DBD, LGF TFs have likely been diverging for 2.5–3 billion years^11,19^. Like their SBP homologs, proteins of the LGF have evolved to recognize a diverse repertoire of molecular effectors in parallel through changes to their effector binding sites, while maintaining allosteric control, suggesting that the capacity to acquire distinct small-molecule ligands evolved multiple times within this family^11,20,21^. Recent phylogenomic analyses reveal a rich diversity of carbohydrate-sensing and global regulators among LacI/GalR TFs, yet questions persist regarding the detailed molecular routes by which novel effector affinities and specificities emerge^22,23^. Studies on related periplasmic binding protein homologs highlight how ancestral sequence reconstruction (ASR) can uncover thermodynamic and structural intermediates that foster promiscuous binding, which then become specialized as evolution proceeds ^4^. Similarly, the use of ASR to probe the evolution of specificity in the DNA-binding domain of the LGF revealed an unexpectedly rugged landscape, which is consistent with the requirement for high fidelity and low promiscuity in the genetic regulation they enable^23^. Applying ASR to the analysis of the evolution of effector specificity in LGF thus has considerable utility in deepening our understanding of the evolution of ligand specificity and advancing our broader understanding of allosteric regulation^24^.

In this study, we have used ASR to reconstruct a trajectory of LGF TFs spanning the last common ancestor (LCA) of the LGF to the extant *Escherichia coli*. LacI. In addition to the finding that binding promiscuity and conformational plasticity are key molecular phenomena in the acquisition of new phenotypes, we observe dramatic thermodynamic trade-offs in entropic and enthalpic binding contributions as a mechanism of adaptive evolution. Specifically, we demonstrate that the most ancient ancestor binds to its cognate ligand, fucose, in an entropically driven mechanism, relying on the hydrophobic effect and retaining similar structure and dynamics to the apo-protein. Through adaptive changes, this binding mechanism radically changes, becoming dominated by specific hydrogen bonding, stabilization of a bound state and significant loss of conformational entropy. We hypothesize that thermodynamic trade-offs in entropy and enthalpy are a common mechanism of evolutionary diversification that extends beyond just LGF TFs, and that entropically-dominated binding modes could have been more common in the distant past owing to the increased thermal energy on Earth in Cambrian/Pre-Cambrian time periods.

## Results

### Phylogenetic reconstruction of the LGF

We have previously reported a reconstructed LGF phylogeny and full ancestral reconstruction methods, although that work focused solely on the DNA-binding domain of ancestral LGF TFs^23^. We reconstructed 10 independent LGF phylogenies by maximum likelihood (ML) using the LG replacement matrix^25^ and a free-rate heterogeneity model^26^, parameterized with 9 discrete rate categories (LG+R9). Of these replicates, none were rejected by the approximately unbiased test^27^ and each tree resolved the same general topology with three major clades and a midpoint placing the LacI lineage as the most ancestral. As all independent trees were statistically indifferent at representing the observed alignment data under the LG+R9 model, a single phylogenetic tree was selected (on the basis of high branch supports at deep nodes in LacI evolution) and used for ML ancestral sequence reconstruction in CodeML of the PAML suite^28^ using the LG replacement matrix with a gamma rate heterogeneity model parameterized with 4 discrete rate categories (LG+G4). The final phylogenetic tree consisted of 519 tips that spanned the full diversity of the LGF (**Figure 1a**)^23^.

**Figure 1.**
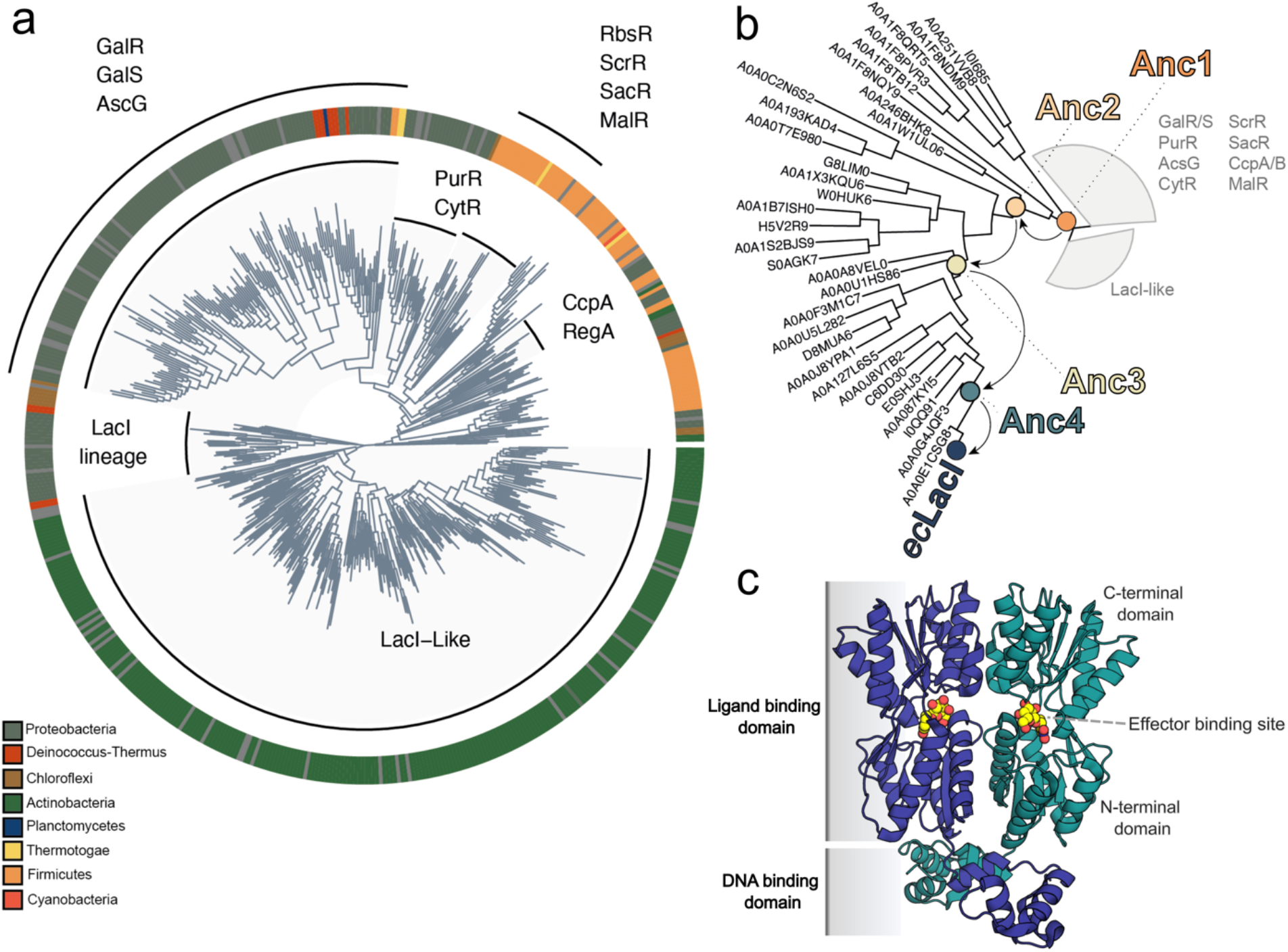
Phylogenetic reconstruction and ASR on the LacI/GalR Family. **a**. Total phylogenetic tree of the LGF **b**. Phylogenetic tree with non-LacI clades collapsed. Nodes belonging to Anc1-4 and ecLacI are highlighted. Filled nodes represent those with branch supports > 90. Nodes belonging to Anc1-4 were each reconstructed with branch supports > 90. **c**. X-ray structure of Lac repressor (PDB ID 2PE5, blue cartoon representation) bound to ligand (ONPG repressor, yellow spheres), crystallized as a canonical dimer (in green and blue cartoons).

Within our dataset, we found extant LGF sequences from diverse bacterial phyla, including *Actinobacteria, Proteobacteria, Deinococcus-thermus, Chloroflexi, Planctomycetes, Thermatogae, Firmicutes* and *Cyanobacteria*. Indeed, the broad taxonomic distribution of the LGF has been described at length previously, leading to hypotheses that the last common ancestor of the family emerged prior to the divergence of the major bacterial lineages^11,29,30^ (~2.5 - 3.2 Gya)^19^. The characterized ancestral sequences were reconstructed with a moderate-high degree of confidence^31^ (mean posterior probabilities ranging from 0.80 - 0.85; **Supplementary Figure 1**); this posterior probability is consistent with the geological timeframe that the LacI superfamily has been diverging over and the consequently sparse phylogenetic signal in extant sequences. Importantly, binding site residues were reconstructed unambiguously and with high posterior probabilities, and we have previously demonstrated that the mean posterior probability of ancestral LacI regulators is not an indicator of function^23,31^. These results together suggest that the ancestral sequences are likely to reconstruct the phenotypes and properties of LGF TFs that emerged, at the earliest, before the last bacterial common ancestor. To investigate the divergence of ligand specificity we selected and experimentally characterized 4 ancestral TFs (Anc1-4) spanning the LCA of the LacI lineage to the extant ecLacI regulator (**Figure 1b, Supplementary Figure 2 & 3**).

### Substrate specificity changes along the LacI evolutionary trajectory

TFs must meet three criteria for functionality: affinity for an effector molecule, the ability to bind DNA and modify gene expression, and allosteric coupling between the effector binding site and the DNA binding domain (**Figure 1c**). We heterologously expressed Anc1-4 and ecLacI in *Escherichia coli* (BL21 DE3) and purified each protein before performing differential scanning fluorimetry (DSF) to screen for binding partners using a diverse library of 192 biologically relevant carbon sources, including sugars, amino acids, amines, amides, and carboxylic acids^32^ (**Supplementary Figure 4**). DSF measures ligand-induced stabilization by measuring the change in melting temperature (T_m_) of a protein in the presence of a ligand. We focused our screening efforts on carbon sources, particularly sugars, as LGF regulators are predominantly involved in carbon metabolism and are typically induced or co-repressed by simple (mono- or di-) carbohydrates and nucleosides. Anc1, Anc3, Anc4 and ecLacI each demonstrated stabilization energies indicative of binding by at least one carbohydrate, with the allolactose analog β-methyl-D-galactoside (BMDG) appearing to stabilize all proteins, other than Anc2. For Anc1, the most ancient ancestor and the LCA of the LacI-lineage, was also significantly stabilized by D- and L-fucose, L-arabinose, and salicin, which was not observed in other ancestors (**Supplementary Figure 5**). BMDG is used as an analog of allolactose, the cognate ligand of ecLacI, due to its enhanced chemical stability that allows for robust and reproducible assay conditions compared to lactose, which has a propensity to hydrolyze under some conditions. Structurally, BMDG consists of an identical D-galactose ring to lactose, with the glucose moiety replaced by a methyl group. These findings are suggestive of an evolutionary transition from binding of lactose/BMDG to D-fucose.

The DSF experiments, along with circular dichroism (CD) measurements, demonstrated that T_m_ values increased towards the most distant ancestor Anc1 (**Supplementary Figure 6**). Indeed, the T_m_ values for Anc1 and Anc2 were measured to be very high (>100 °C and 87.4 °C, respectively; **Figure 2a**). This provided a possible explanation for the lack of observable stabilization of Anc2 in the DSF screen, where ligand binding might not cause a detectable increase in T_m_ of a highly thermostable protein. The high thermostability of these ancestors is consistent with other ASR studies^33^, and in the context of the ambient temperature when they have been predicted to exist (2.5–3.5 billion years ago); a hot ancient climate, where high thermostability would have been critical^34^. This initial screen of 192 potential ligands established that, as for the wild-type ecLacI, the ancestors are relatively specific, binding a small subset of carbohydrate molecules with high specificity.

**Figure 2.**
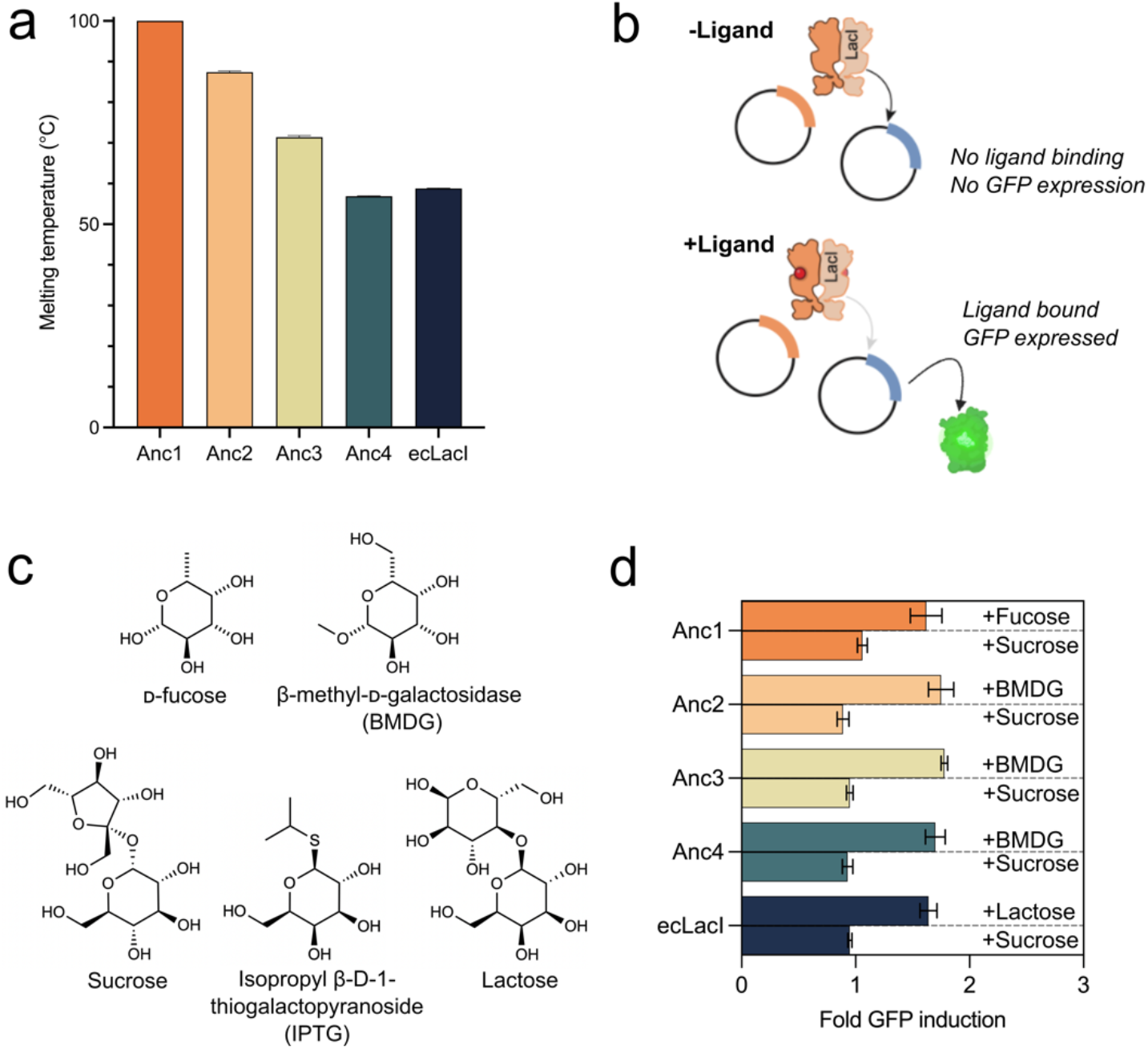
**a**. Melting temperatures (T_m_) of ancestral TFs from DSF experiments, with SEM shown as error bars (n=3). **b**. Illustration of the *in vivo* assay selection system, a dual-plasmid reporter system, where GFP is repressed by the ancestral TF until ligand binding occurs. **c**. Structures of D-fucose, BMDG, IPTG, and sugars used in flow cytometry experiments as positive control (lactose) and negative control (sucrose). **d**. *In vivo* FACS results comparing average GFP fluorescence for ligand vs sucrose negative control, normalized to no ligand. SEM shown as error bars (n=3). Complete results in **Supplementary Figure 8**.

Following our identification of potential ligands, we then tested whether the TFs could respond to their identified ligands *in vivo* to regulate gene expression, i.e. whether ligand binding could allosterically affect the binding of the DNA binding domain to the operator sequence. To achieve this, we used a dual plasmid reported system in which GFP is constituently expressed under control of a *lac* operator (*lacOsym*, a synthetic operator that allows for tighter repression and more reproducible results)^35^. In this system, GFP expression is repressed by a functional TF until binding of a ligand allosterically decreases the affinity of the DNA-binding domain and operator, allowing GFP to be expressed and measured using flow cytometry (**Figure 2b, Supplementary Figure 7**). To avoid interference by endogenous LacI and other carbohydrate sources, we used

*E. coli* 3.300 cells, which have an interrupted *lacI* gene (thus no native LacI protein), grown in minimal media^36^. Cells were transformed with plasmids encoding the reporter system and tested with sucrose as a negative control that reported essentially same response as when no ligand was added, i.e. no increase in GFP expression, BMDG, D-fucose and lactose (positive control) (**Figure 2c**). As expected, when we express wild-type ecLacI in this system, very low levels of GFP are expressed until lactose is added, where a 65% increase in GFP is reported. We observe similar results for the ancestral TFs, where the ancestral TFs act as repressor proteins *in vivo*, and specific ligands (matching those predicted from the DSF screen) induce an allosteric response that increases GFP expression. LacI and Anc2–4 exhibit strong responses to lactose and BMDG, while D-fucose is the strongest inducer of Anc1, suggesting that ligand specificity seen *in vitro* is translated to allosteric transcriptional regulation *in vivo*. (**Figure 2d, Supplementary Figures 8–10**).

### The thermodynamics of binding is an evolvable property

To probe the thermodynamic parameters that have shaped ligand specificity in LGF TFs across evolution, we performed isothermal titration calorimetry (ITC). Having established that our ancestral TFs are indeed capable of phenotypic regulation in response to BMDG and D-fucose, we used ITC to elucidate the dissociation constant (*K*_D_) and the enthalpic (ΔH) and entropic (-TΔS) contributions to the free-energy of binding (ΔG_binding_), that each regulator exhibited towards its various ligands.

The affinity across Anc1 to ecLacI for BMDG increased over approximately an order of magnitude along the evolutionary trajectory; from 4 mM for Anc1, to 2.2 mM for Anc2, 2.5 mM for Anc3, 550 μM for Anc4 and 306 μM for ecLacI (**Figure 3a, Supplementary Table 1**). Given that BMDG is a known inducer of ecLacI and a structural analog of lactose, this is not surprising and consistent with selective pressure to bind to the cognate galactose moiety. In contrast, D-fucose displayed high affinity to the most ancient ancestor, Anc1 (*K*_D_ of 127 μM), with little to no affinity observed in the latter ancestors or ecLacI; the only variants to display detectable binding were in Anc4 and ecLacI, which exhibited *K*_D_ values of 15 mM and 37 mM, respectively, approximately 100-fold lower than Anc1 (**Figure 3b, Supplementary Table 1**). The *K*_D_ of Anc1 for D-fucose is similar in magnitude to that of typical extant proteins of the LacI/GalR superfamily for their cognate ligands, e.g. GalR has a reported *K*_D_ for D-fucose of 60 μM^37^. This result suggests that the LCA of the ecLacI lineage likely bound to simple pyranose-base molecules, such as D-fucose, but had some promiscuous affinity for galactose/lactose, which was then selected for through evolution to yield the specialized extant ecLacI protein. The diversification of these ancient LacI ancestors to recognize β-substituted D-galactosides (such as BMDG and allolactose) probably occurred as early prokaryotic metabolism became more complex, requiring more extensive regulation. Indeed, D-fucose (among other simple pyranoses) likely emerged as a primary carbon source earlier than β-substituted galactosides such as lactose and allolactose (1–2 Gya *vs*. 3.5 Gya)^38^. The non-productive (no transcriptional regulation in the *in vivo* assays) and low-affinity binding that Anc4 and ecLacI display towards D-fucose are thus likely to be vestigial properties that are retained from the ancestral preference for D-fucose. Transcriptional regulation is of utmost importance to more complex life, hence, the divergence and radiation of the LacI/GalR family tree may represent evolving bacterial complexity.

**Figure 3.**
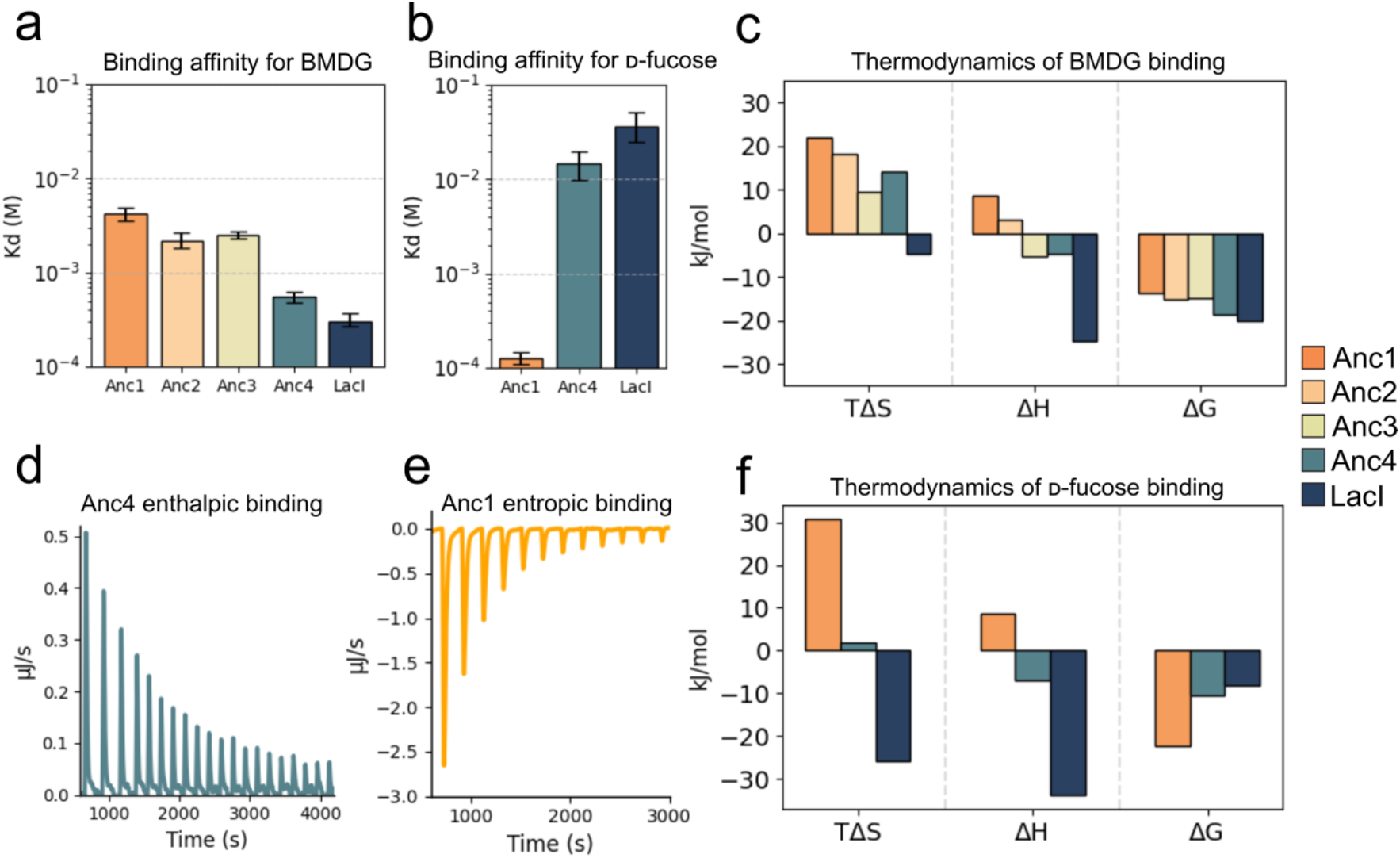
Isothermal titration calorimetry was used to probe the binding affinities (K_D_) and thermodynamics binding parameters of the ancestral transcription factors **a**. Binding affinity (M) of ancestors for BMDG and **b**. D-fucose with error bars of 68.3% CIs from at least n=3 titrations. **c**. Thermodynamic parameters for binding of Anc1-4 and LacI to BMDG from global analysis. **d. & e**. Contrasting entropic and enthalpic modes of binding for Anc1:D-fucose and Anc4:BMDG, illustrated with indicative thermograms, full thermograms in **Supplementary Figure 11. f**. Thermodynamic parameters for binding of Anc1, Anc4, and LacI to D-fucose from global analysis. Full ITC data in **Supplementary Table 1**.

Enthalpy-entropy trade-offs in the adaptive evolution of these proteins have been documented previously in the broader SBP superfamily to which the ligand binding domains of the LacI/GalR family belong, where promiscuous binding of two ancestral SBPs to amino acids exhibited different thermodynamic signatures^4^. On examination of the thermodynamic parameters that contribute to the binding affinity of the ancestral transcription factors, the ΔG_binding_ of BMDG decreased from −13.6 kJ/mol for Anc1 to −20.1 kJ/mol in ecLacI, which was the result of more extensive changes in the thermodynamic contributions of enthalpy and entropy. ITC revealed that Anc1 and Anc2 bind BMDG *via* an entropically favored (TΔS = 22.1 kJ/mol and 18.2 kJ/mol; ΔH = 8.53 kJ/mol and 3.01 kJ/mol, respectively) endothermic mechanism. In contrast, extant ecLacI was found to bind BMDG via an exothermic and enthalpically favored mechanism (TΔS = −4.9 kJ/mol, ΔH = −24.9 kJ/mol) (**3c & d, Supplementary Table 1**). The evolutionarily intermediate species Anc2 and Anc3 are also intermediate in terms of their binding thermodynamics, displaying relatively balanced thermodynamic profiles where binding is both entropically and enthalpically favorable, albeit with smaller contributions from each thermodynamic property such that their sum is comparable to other regulators at ambient temperature. Importantly, similar thermodynamic trends were observed when D-fucose was tested with Anc1, Anc4 and ecLacI: the endothermic and entropically driven binding of D-fucose in Anc1 becomes exothermic and enthalpically driven in Anc4/ecLacI (**Figure 3e & f**). It is notable that the change in binding thermodynamics appears to be a gradual trade-off spanning the trajectory, suggesting that the thermodynamic driving force of a binding event is a molecular property under evolutionary pressure rather than an artefact of the reconstruction.

### The structural basis for entropically *vs*. enthalpically driven ligand binding

To understand the molecular basis for the observed enthalpy-entropy trade-offs, we solved the structures of Anc1 and Anc4 in complex with their ligands by X-ray protein crystallography (**Supplementary Table 2**). As we focused exclusively on small-molecule binding in the ligand binding domain (LBD), we truncated the 60 N-terminal amino acids comprising the DNA binding domain (Δ). To confirm that this would not affect the ligand binding, we repeated all ITC experiments on each ΔAnc variant and observed the same binding trends as with full-length regulators (**Supplementary Table 3, Supplementary Figure 12 & 13**), confirming that structural studies limited to the LBD can provide insight into small-molecule recognition of the full TFs.

We solved the crystal structure of ΔAnc4 in a complex with BMDG at a resolution of 1.50 Å. The ΔAnc4:BMDG complex crystallized as a canonical LGF dimer in the asymmetric unit. BMDG was bound with full occupancy and interacts with ΔAnc4 *via* hydrogen bonding from the BMDG methoxy substituents to T69 and D148 and the BMDG 2’ and 3’ hydroxyl moieties to R195, N243, D271 and Q288 (**Figure 4a**). These residues are conserved with ecLacI and make homologous contacts to IPTG (which is identical to BMDG in all respects except the BMDG methoxy group is replaced by sulfur-linked isopropyl group, **Figure 2b**) in extant ecLacI (via S69, D149, R197, N246, D274, Q291; **Supplementary Figure 14**). Indeed, when ΔAnc4 and ecLacI are superimposed, there are a number of similarities, such as the effector binding location, orientation, and contacts, and the closed conformation of the bound structure. A molecular dynamics simulation of ΔAnc4 without BMDG in the binding site (starting from the closed ligand bound conformation) showed relaxation to an open *apo* structure within 100 ns for four replicates, indicating that Anc4 likely undergoes an open-closed change during ligand binding, analogous to ecLacI (**Figure 4b, Supplementary Figure 15**). EcLacI has a greater enthalpic contribution to binding of BMDG. This can be rationalized by the sequence differences T69S, V126Y, P161H (H163 in ecLacI) and T188S (S190 in ecLacI) which introduce additional hydrogen bonding interactions into LacI. Thus, the enthalpic binding mode observed in Anc4 with BMDG appears relatively straightforward and consistent with our understanding of ecLacI^39,40^.

**Figure 4.**
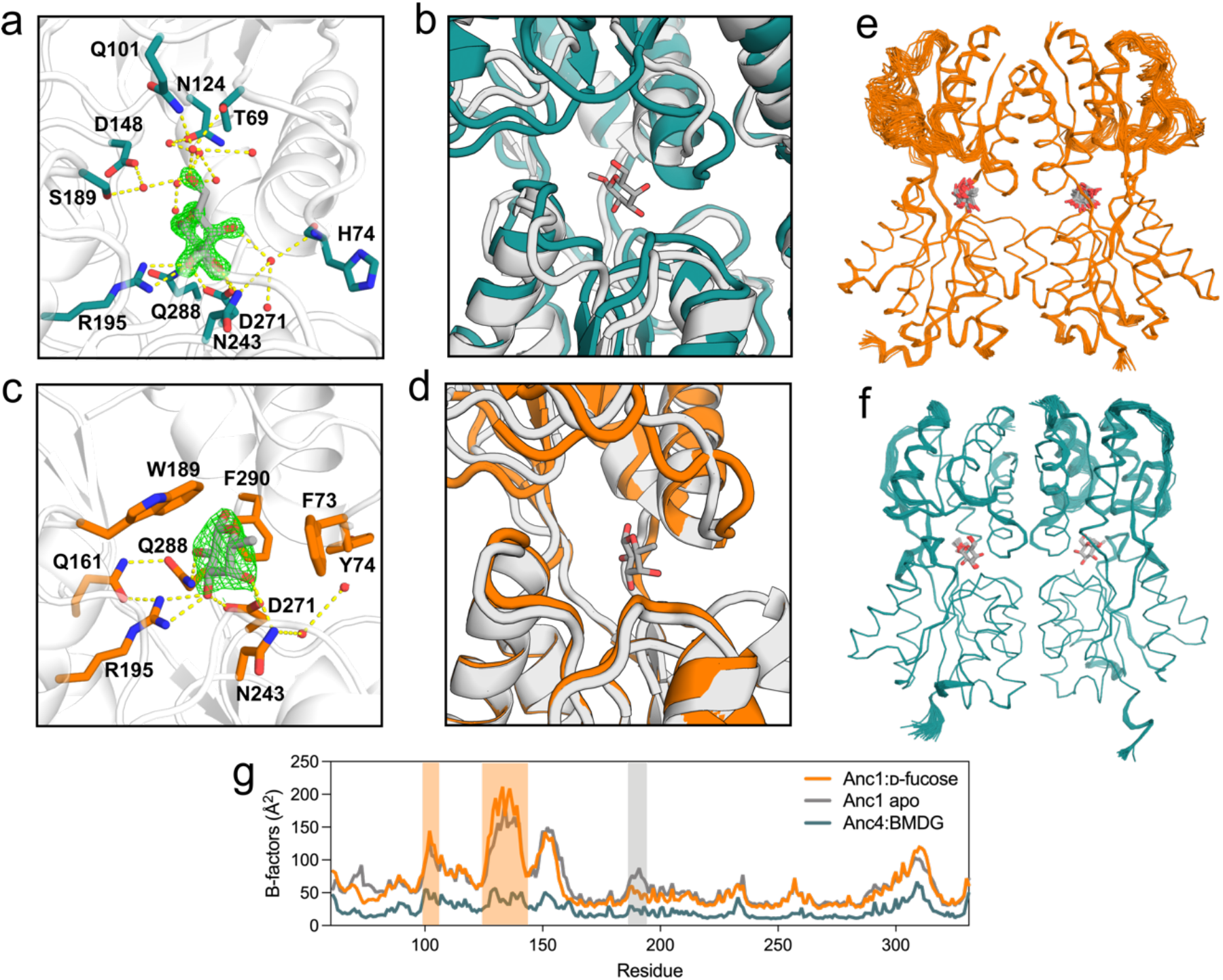
Binding pockets of ancestral LGF members with omit electron density (F_o_–F_c_) shown in green mesh contoured to 2σ, waters within 4 Å shown as red spheres. **a**. ΔAnc1 bound to D-fucose. W218 excluded for clarity. **b**. ΔAnc4 bound to BMDG. **c**. ΔAnc1:D-fucose complex binds in open conformation (orange) overlay with ΔAnc1 apo (white). **d**. ΔAnc4:BMDG complex binds in closed conformation (green) overlay with ΔAnc4 apo from relaxation MD simulations (white). Ensemble refinement of **e**. ΔAnc1 (D-fucose shown in grey) and **f**. ΔAnc4 (BMDG shown in grey), (**Supplementary Table 5** for ensemble refinement statistics) **g**. Average B-factors (Å^2^) for each TF plotted by residue, with areas of interest in ΔAnc1 shaded (in orange, loop T98–D104 & helix A125–P143, and in grey, loop G186–R193).

To better understand the entropically favorable binding of D-fucose by Anc1, we obtained a complex of ΔAnc1 with D-fucose **(Figure 4c**). The ΔAnc1:D-fucose complex diffracted to 2.5 Å (**Supplementary Table 2**), with eight monomers in the asymmetric unit, organized as four LacI-like dimers (**Supplementary Figure 16**). D-fucose was observed to be bound in the active site of all 8 monomers at full occupancy. Interestingly, the ligand bound to a mostly open state of the LBD (**Figure 4d**) and is coordinated by hydrogen bonds from the 2’ and 3’ hydroxyl groups on D-fucose to R195, N243, D271, and Q288, residues that make analogous contacts in BMDG. ΔAnc1 shows several hydrophobic contacts, F73, W189, W218, F290, where F73 and W189 are unique to ΔAnc1. Analysis of the binding pocket of ΔAnc1 shows only two coordinating crystallographic waters that interact with the 4’ hydroxyl group and mediate a hydrogen bond interaction to the main chain amide of Y74 (**Supplementary Figure 17**). The importance of this site was probed by mutation to the ΔAnc4 H74 residue, resulting in loss of binding to both BMDG and D-fucose

(**Supplementary Figure 18**). This is likely due to a conformational change seen in H74, that would have disrupted this important water-mediated hydrogen bond interaction between D-fucose and the ΔAnc1-H74 mutant.

The water networks in the Anc4:BMDG complex are vastly different to ΔAnc1, where a network of coordinating crystallographic water molecules can be seen in the binding pocket; BMDG contacts the N-terminal subdomain of ΔAnc4 *via* two ordered water molecules that bridge T69, H74 and Q101 and to the C-terminal domain *via* an additional two waters molecules that contact N124, D148 and S189. These waters themselves make further bonds to ordered waters, to a total of 12 ordered crystal waters in the ΔAnc4 binding pocket (**Figure 4a, Supplementary Figure 19**). Examination of the binding pocket regions of ΔAnc1 and ΔAnc4 show mutations at the sites F73L, Y74H, G75A, S148D, Q161P, W189S and E191S (**Supplementary Figure 20**). Generally, the binding pocket of ΔAnc1 is characterized by bulky aromatic residues that evolved to smaller, more polar residues in ΔAnc4. However, some mutations in the evolutionary trajectory from ΔAnc1 to ΔAnc4 involve an increase in hydrophobicity or loss of charge (i.e., Q161P, E191S) and highlights the complex interactions in ligand binding that make rational design challenging. It was also notable that the structure of the ΔAnc1:D-fucose complex was considerably less well ordered than the ΔAnc4:BMDG complex (**Figure 4e & f**), with loop E115–G120, helix N128–V141, and loop F146–T156 difficult to resolve in the electron density. The eight D-fucose molecules bound to ΔAnc1 appear to retain some level of mobility even when bound in the binding pocket; the average B-factors range from 41.4–56.9 Å^2^ (**Supplementary Table 4**). This is in comparison to the much lower B-factors of the two BMDG ligands found in dimeric ΔAnc4, of 23.6 and 21.0 Å^2^. In summary, the binding of D-fucose to Anc1 is consistent with an entropically favorable mechanism in which the coordination is considerably less reliant on electrostatic interactions (H-bonds) than in the comparative ligand bound structures of Anc4 and ecLacI, and the structure retains considerable levels of disorder, even when bound to its ligand.

To understand the entropically favorable binding of D-fucose by ΔAnc1 in more detail, we also solved the structure of the apo protein at high (1.67 Å) and moderate (2.3 Å) resolution (**Supplementary Table 2**). The high-resolution structure had glycerol bound in the active site (**Supplementary Figure 21a**), whereas the moderate resolution structure had only water molecules present in the active site. These structures allowed us to make some clear observations: first, ΔAnc1 is intrinsically very disordered; even the high-resolution structure was difficult to model owing to the mobility of the small N-terminal domain (**Figure 4g**). Secondly, the apo-structure of ΔAnc1 in the open state was virtually indistinguishable from the ligand bound structure in the binding site i.e., there was almost no loss of disorder upon ligand binding, which is consistent with the low entropic penalty, while a number of water molecules are observed to be liberated by ligand binding to make binding entropically favorable (**Figure 4c**). Together, these results provide a molecule-level understanding of the entropically favorable ligand binding mode observed in Anc1.

### The evolution of ligand specificity in ancestral TFs

Analysis of the crystal structures of ΔAnc1:D-fucose and ΔAnc4:BMDG allows us to understand both ligand specificity and the thermodynamics that drive ligand binding. Comparison of the binding pockets of ΔAnc1 and ΔAnc4 show that D-fucose binds in a different position to BMDG, and thus also to IPTG in ecLacI, with D-fucose binding deeper in the binding pocket to accommodate for the presence of W189 (**Figure 4a & b, Supplementary Figure 20**). It is likely that W189 prevents BMDG binding by sterically clashing with the galactoside 6’ hydroxyl group that is absent in D-fucose. Examining the evolutionary trajectory from ΔAnc1 to ΔAnc4, it appears that evolution favors removal of bulky aromatic/hydrophobic groups such as F73 and W189, instead replacing them with residues capable of forming direct or water mediated hydrogen-bonds in ΔAnc4. We hypothesize that, upon binding of D-fucose, the desolvation of water molecules from these hydrophobic residues in ΔAnc1 would increase the favorable entropic contribution of the hydrophobic effect.

Anc1 is distinguished by its ability to bind D-fucose in an open state, while Anc4 binds BMDG in a more closed conformation (**Figure 4c & d**). Along the evolutionary trajectory, the closed state is enriched where it becomes the functional ligand binding confirmation in Anc4. This change is enabled by structured water networks that bridge BMDG to the N-terminal domain in ecLacI and Anc4 that are disrupted in Anc1 due to ancestral W189 and F73. This effect explains the difference in the openness of binding pockets in the two ancestors; D-fucose lacks bridging contacts between the two domains in Anc1, while BMDG contacts both domains, closing the two domains together. As we observed *in vivo*, Anc1 is still able to impart allosteric control over DNA transcription, despite not undergoing a large open-closed conformational change in response to ligand binding, as is the understood mechanism of TFs. However, Anc1 does undergo large conformational shifts distal to the binding site (loops T98–D104, G186–193, helix A125–P143) when compared to the apo structure (**Figure 5a, 4g**), that we propose are sufficient for the continued allosteric functionality of Anc1. Flexibility and promiscuity are common traits in ancestral proteins, likely reflecting their role as evolutionary starting points with broad binding potential. In Anc1, this flexibility not only supports future evolvability but also contributes to its strongly entropically-driven binding.

**Figure 5.**
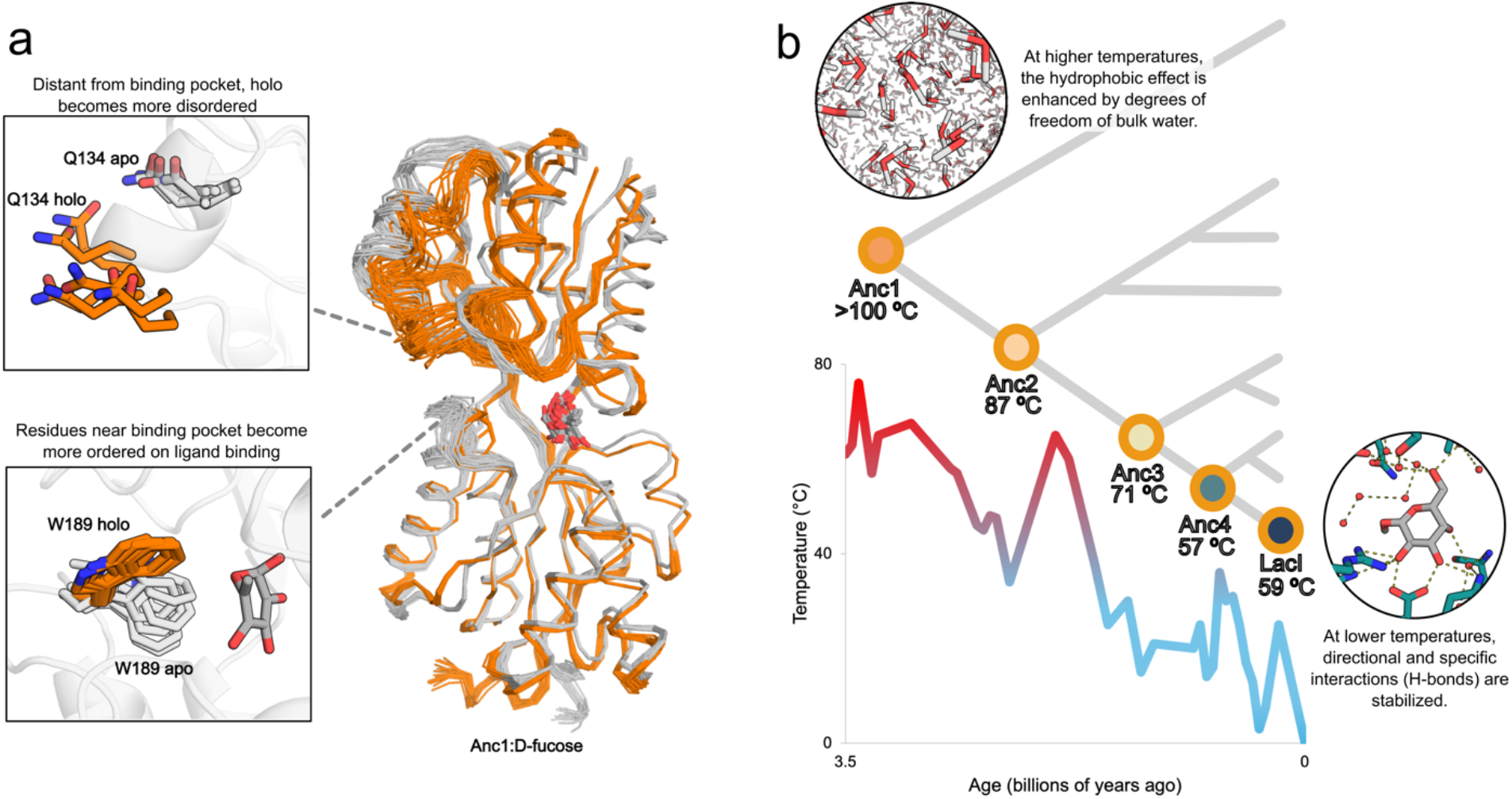
**a**. Overlay of ensemble refinement of Anc1 apo (grey) and holo (orange). Entropic binding in Anc1 may be offset by redistribution of disorder from the binding pocket to distal regions on binding of D-fucose; for example, analysis by ensemble refinement shows that W189 holo (orange) has fewer degrees of freedom on binding compared to the apo structure (grey), while Q134 becomes more dynamic after binding. **b**. Simplified phylogenetic tree of the LGF, shown with melting temperatures of each ancestor calculated using DSF, that correlate with increase in ancient environmental temperatures (shown as ocean temperatures)^34^. Timescales for ancestors are estimates.

The favorable entropy of binding of Anc1 to D-fucose is amplified by its retention of disorder when bound. This concept, termed spatial entropic redistribution, describes how loss of entropy in binding site residues can be compensated by redistribution of entropic freedom to distal regions of the protein structure, reducing the overall free energy lost on binding (**Figure 5a**)^41,42^. While binding is generally entropically unfavorable due to loss of conformational entropy in both the protein and ligand^43^, the hydrophobic effect (release of water molecules), and Anc1’s ability to bind D-fucose with retained disorder in the N-terminal domain, offset this penalty. It should be noted that he apo and D-fucose-bound ΔAnc1 structures were obtain from crystals from same crystal screening drop i.e., the observed differences are not due to differences in crystallization conditions.

The trend in binding thermodynamics, and the thermostability of these ancestral proteins, are consistent with the hypothesis that the LCA of the ecLacI clade likely emerged 2.5 - 3.2 Gya^19,44^ in a hot geobiochemical climate^34,45–47^. DSF and CD experiments demonstrated that T_m_ increased with each ancestor, where Anc1 was observed to have T_m_ > 100 ºC (**Figure 2a, Supplementary Figure 6**). Likewise, the entropic contribution to free energy is modulated by temperature through the −TΔS term; under high-temperature conditions, such as those in a Precambrian climate, the amplified contribution of the entropic term would favor entropically driven binding. As global temperatures decreased, the relative advantage of a large TΔS contribution would diminish, potentially shifting selective pressures toward binding interactions where the free energy is dominated by enthalpic factors (**Figure 5b**), while the thermostability of the proteins would also decrease (**Figure 2a**). These observations suggest that molecular binding mechanisms have evolved, at least in part, in response to the prevailing environmental temperatures, with early thermophilic organisms developing traits optimized for stability and binding efficiency at high temperatures.

## Discussion

In this work, we have shown that LGF TFs evolved distinct ligand specificity via entropy-enthalpy trade-offs. One of the most notable aspects of the LGF evolutionary trajectory is the gradual change in thermodynamic binding parameters from most distant to most recent ancestor, which suggests thermodynamic trade-offs in entropy and enthalpy of effector binding are a central part of adaptive evolution. Along the evolutionary trajectory, the binding is switched from entropically-driven in the most distant Anc1, to enthalpically-driven in most recent Anc4 and ecLacI. This trade-off is accompanied by a switch in specificity from D-fucose in Anc1 to BMDG in ecLacI and recent ancestors, where high affinity binding of Anc1 to D-fucose could reflect the metabolic relevance of D-fucose in the ancestral organism’s environment.

Structural analysis using X-ray crystallography shows that entropically driven binding in Anc1 is accommodated by an open ligand binding pose, the retention of conformational disorder through entropic reservoirs, and contributions from the hydrophobic effect. D-fucose retains degrees of freedom in the Anc1 binding pocket, contrary to the idea that ligands lose their conformational entropy on binding^48^. The hydrophobic binding pocket residues in Anc1 evolve to smaller, polar residues in Anc4 that result in the formation of a number of enthalpically favorable, but entropically unfavorable, water-mediated hydrogen bonds between BMDG and Anc4/ecLacI. Thus, we observe clear molecular-level explanations for the gradual shift from entropically-to enthalpically-driven ligand binding in this evolutionary trajectory.

Our analysis reveals that thermodynamic binding mechanisms in ancestral transcription factors likely evolved in response to changing global temperatures, with early proteins favoring entropically driven binding modes that leverage retained structural disorder and water release to offset the conformational entropy cost of ligand binding, exemplified by Anc1. As environmental temperatures decreased over evolutionary time^34,45–47^, the selective advantage of large −TΔS contributions diminished, and binding mechanisms gradually shifted toward more enthalpically dominated interactions, as seen in Anc4 and ecLacI. This transition coincides with a structural tightening of the ligand-binding pocket, increased hydrogen bonding, and the emergence of structured water networks that stabilize the ligand-bound state. These changes are also fully consistent with the observed trend in thermostability. Thus, the evolution of binding thermodynamics reflects not only ligand specificity and structural constraints, but also adaptation to Earth’s cooling climate, highlighting temperature as a key, and under-appreciated, selective force shaping molecular recognition.

Altogether, these results suggest that diversification of molecular function in the LacI/GalR family occurred through adaptive thermodynamic trade-offs. By reconstructing and characterizing ancestral transcription factors, we uncovered a continuum from entropy-driven binding in ancient thermophilic ancestors like Anc1 to enthalpy-dominated binding in more recent ancestors and extant LacI. Our structural analyses revealed how these shifts were mediated by changes in conformational dynamics, water networks, and binding pocket composition, offering a detailed molecular rationale for altered ligand specificity and thermodynamic strategy. This study underscores the value of ASR; this level of mechanistic insight is arguably only possible through ancestral sequence reconstruction, which uniquely enables the study of long evolutionary trajectories revealing how complex properties like thermodynamic binding strategies emerge and shift over billions of years. These findings demonstrate that the evolution of protein function involves not only changes in sequence or structure but also rebalancing the energetic components of ligand binding in response to environmental pressures such as global temperature shifts. More broadly, this work highlights the potential for enthalpy–entropy compensation to serve as a general mechanism by which new biochemical phenotypes emerge over evolutionary time.

## Methods

### Phylogenetic analysis and ancestral sequence reconstruction

See Meger et.al (2024) for previously described comprehensive LGF phylogenetic reconstruction methods ^23^.

### Cloning and mutagenesis

Ancestral sequences were reverse translated and codon optimized for translation in *E. coli*. Synthetic genes for the ancestral protein were synthesized, cloned into the pET28a(+) vector and sequenced by Twist bioscience. The genes were cloned such that each included a C-terminal TEV cleavage site preceding a hexahistidine tag. The C-terminal was chosen as the location for the histidine tag as the N-terminal is close to the DNA binding domain, whereas the C-terminal is relatively distant from known functional sections of LGF proteins yet is still solvent exposed. To keep each gene in-frame with the vector encoded start codon and C-terminal tag, the ancestors contained an additional methionine and glycine at their N-termini. The coding sequence for the *E. coli* lac repressor was codon optimized for recombinant expression and synthesized with 3’ and 5’ homologous overlap with the empty pETMCSIII multiple cloning site, a hexahistidine tag and a TEV cleavage site. The linear LacI DNA was cloned into pETMCSIII by Gibson assembly under the T7 promoter.

### Protein expression

Plasmids containing genes for the maximum likelihood ancestral reconstruction of nodes 1142 (Anc4), 1136 (Anc3), 1133 (Anc2) and 1129 (Anc1) were obtained from Twist Bioscience. The pET28a(+) vector with C-terminal His6 tag after a TEV cleavage site was used. Proteins were expressed in *E. coli* (BL21)DE3 cells and grown in Luria-Bertani (LB) supplemented with 100 mg/L kanamycin (Sigma). Cultures were grown to OD_600_ 0.6 at 37 ºC, induced with 1 mM β-d-1-isopropylthiogalactopyranoside (IPTG) and incubated for a further 18 hours at 20 ºC, 180 rpm. Cells were pelleted and stored at −20 ºC before purification.

### Protein purification

Proteins were purified under native conditions by nickel-nitrilotriacetic acid (Ni-NTA) affinity chromatography and size-exclusion chromatography (SEC). Cells were thawed, resuspended in equilibration buffer (20 mM Tris-HCl, 500 mM NaCl, 20 mM imidazole, 2% glycerol, pH 8.0) and lysed with sonication and addition of 0.5 μL turbonuclease. Soluble and insoluble fractions were separated by centrifugation (21000 x *g* for 30 min at 4 ºC). The supernatant was filtered using a 0.2 μm filter and loaded onto a pre-equilibrated 5 mL HisTrap HP column (GE Healthcare). Hexahistidine-tagged protein was eluted with an elution buffer (20 mM Tris-HCl, 500 mM NaCl, 250 mM imidazole, 2% glycerol, pH 8.0). For crystallography and ITC, protein was exchanged into TEV protease-cleavage buffer (50 mM Tris-HCl, 150 mM NaCl, 1 mM DTT, 0.5 mM EDTA, pH 8) using a pre-equilibrated desalting column (GE Healthcare). TEV cleavage with 1:10 ratio of TEV protease to protein was performed for 18 hours at 25 ºC for ΔAnc1-3, while ΔAnc4 was incubated at 4 ºC. Cleaved protein was passed over a pre-equilibrated 5 mL HisTrap HP column (GE Healthcare) to remove cleaved hexahistidine tags. Protein was concentrated and using a centrifuge filter (Amicon Ultra-15 filter unit with 10 kDa cut-off, Merck Millipore) and purified by SEC on a HiLoad 26/600 Superdex 200 column (GE Healthcare), typically eluting in SEC buffer (20 mM HEPES, 150 mM NaCl, 2% glycerol, pH 8). Protein purity was confirmed by SDS–PAGE, and protein concentrations were measured spectrophotometrically (NanoDrop Technologies) using molar absorption coefficients calculated in ProtParam.

### Crystallization and structure determination

*ΔAnc1:glycerol*. Protein was concentrated to 22 mg/mL in SEC buffer using 10 kDa MWCO centrifugal concentrator prior to crystallization. Crystals were obtained within 10 days using the hanging drop vapor diffusion method against a reservoir solution containing 14.4% PEG 8000, 0.08 M sodium cacodylate pH 6.2, 0.16 M calcium acetate, 20 % glycerol. The crystallization drop was comprised of 2 μL protein and 2 μL well solution. Crystals were flash cooled in liquid nitrogen without further cryoprotecting.

*ΔAnc1 apo*. Protein was concentrated to 40 mg/mL in SEC buffer using 10 kDa MWCO centrifugal concentrator prior to crystallization. Crystals were obtained within 1 days using the hanging drop vapor diffusion method against a reservoir solution containing 15%w/v PEG 4000, 0.2 M MgCl_2_, 0.1 M Tris-HCl, pH 8.5 The crystallization drop was comprised of 1 μL protein and 1 μL well solution. Crystals were flash cooled in liquid nitrogen without further cryoprotecting.

*ΔAnc1:D-fucose*. Apo crystals from above were soaked by adding 2 μL of 200 mM D-fucose dissolved in water to the apo crystal drop and left to incubate up to 1 minute. Crystals were flash cooled in liquid nitrogen without further cryoprotecting.

*ΔAnc4:BMDG*. Protein was concentrated to 22 mg/mL in SEC buffer using 10 kDa MWCO centrifugal concentrator prior to crystallization. Concentrated protein was incubated for 30 minutes at RT with 3.6 mM BMDG dissolved in water prior to crystallisation. Co-crystals were obtained within 2 days using the hanging drop vapor diffusion method against a reservoir solution containing 21 % PEG 3350, 0.1M HEPES pH 7.1. The crystallization drop was comprised of 2 μL protein and 2 μL well solution. Crystals were flash cooled in liquid nitrogen with cryoprotectant (35% PEG 3350, 0.1 M HEPES pH 7.3, 3.6 mM BMDG).

Diffraction data were collected at 100 K on the MX2 beamline of the Australian Synchrotron ^49^. Datasets consisting of 3600 images were collected (1 s/degree exposures) on a Dectris EIGER 16M detector using a wavelength of 0.95372 Å. Diffraction data were processed using XDS and AIMLESS ^50,51^ and phased by molecular replacement using Phaser ^52^ from the CCP4 suite ^51^, and using the lactose repressor bound to IPTG (PBD: 2P9H) structure as the search model for ΔAnc4:BMDG and ΔAnc1:glycerol (sequence similarity with ΔAnc1 38.3%, ΔAnc4 similarity 72.9%) and AF2 models ^53^ for ΔAnc1:D-fucose and apo ΔAnc1. Refinement was conducted using REFMAC ^54^ and Phenix ^55^, and model building with Coot ^56^ and ISOLDE ^57^. Structures have deposited to the Protein Data Bank under IDs 9NT3, 9NT4, 9NT5, 9NT6.

Full details of crystallization and structure determination for each protein are given in **Supplementary Tables 2**. Data collection and refinement statistics are given in **Supplementary Table 2**. Representative electron density for the active/binding site of each structure is given in **Supplementary Fig. 19**.

### Differential scanning fluorimetry

DSF experiments were performed with 0.5 mg/mL protein and using the thermal shift dye (Thermofisher) and a Quant Studio3 Real-time PCR thermocycler (ThermoFisher). The library of small molecules was bought commercially from Biolog (Microbial phenotyping plates PM1 and PM2A). The concentration of each molecule in the library was between 10 and 20 mM. Master mix, consisting of protein in buffer with the fluorescent thermal shift dye, was assayed against each well in the small molecule library. Melting temperatures were computed using the ThermoFisher Thermal-Shift analysis software suite as the first derivative of fluorescence. DSF experiments were performed in duplicate.

### Circular dichroism spectroscopy

Spectra was attained using an Applied Photophysics Chirascan CD spectrophotometer, using a 1 mm quartz cuvette. Each protein was diluted to 0.2 mg/mL in MQ water. An initial background scan using a buffer-only blank was subtracted before spectra acquisition between 180 to 280 nm, using a bandwidth of 1, step size of 0.5, and temperature at 20 ºC. Each run consisted of three traces and was duplicated. Melting temperatures (T_m_) were obtained by fitting a sigmoidal curve to data at 208 and 222 nm respectively from 20 ºC to 90 ºC.

### Isothermal titration calorimetry

Purified ancestors and ligands were prepared in the same HEPES buffer used for SEC purification and filtered and degassed, and ligand concentrations prepared fresh for each titration. A TA Instruments benchtop Nano-ITC (small volume) was used to perform the titrations. Proteins were concentrated using an Amicon Ultra-15 filter unit with 10 kDa cut-off (Merck Millipore). Ligand was titrated into protein, starting with a 1 μL dummy injection, then 22 2 μL injections, with a 250 s gap between titrations. Experiments were conducted at 25 ºC, with a stirring of 300 rpm. Initial and final baselines were generated over 400 s. Analysis of results was conducted first using NITPICK ^58^ to find the baseline, then SEDPHAT ^59^; the baseline-subtracted power was integrated, and the integrated heats fitted to the single-binding site model (A + B ↔ AB, heteroassociation) to obtain Ka and ΔH. Values for the incompetent fraction of protein were constrained between −0.2 and 0.2. Global fitting was obtained by iterative cycling between Marquardt-Levenberg and Simplex algorithms in SEDPHAT ^59^ until convergence of model parameters was observed. 68.3% confidence intervals were calculated using the automatic confidence interval search in SEDPHAT ^59^. Thermograms were plotted using Gussi ^60^.

### Intrinsic tryptophan fluorescence

Ancestral mutant Anc1-Y74H was made to final concentration of 10 μM in relevant concentrations of D-fucose or BMDG, centrifuged to mix, and incubated for 10 minutes at RT. Tryptophan fluorescence was then measured using a TECAN infinite M200 plate reader with excitation fixed at 260 nm and emission spectra collected between 288–400 nm.

### Ensemble refinement

Time-averaged ensembles were generated for Anc1 and Anc4 with phenix.ensemble_refinement implemented in PHENIX version 1.21. Prior to ensemble refinement, the ancestral structures were refined using phenix.refine with default parameters (the non-bonded crystallographic condition ligands EGO PEG and ACT were removed). The R_free_ of the refined models, and the mFO − DFC difference density maps, were used to assess the time-averaged refinement protocol. To prepare the structures for ensemble refinement, alternate conformations were removed and occupancies adjusted to 100% (PDB tools). Hydrogen atoms were added (phenix.readyset). The pTLS parameters were optimized using phenix.refine to minimize the Rfree value by systematically testing different partitioning of the model into TLS (Translation/Libration/Screw) groups; full ensemble refinement statistics are detailed in **Supplementary Table 5**.

### Molecular dynamics simulations

MD simulations of Anc4 were initialized from the BMDG bound structure that had the ligand removed. Simulations were performed using GROMACS version 2024.1, using the CHARMM36m forcefield ^61^. GROMACS was used to add terminal charged amide and carboxylic acid groups, based on simulation conditions of pH 7.5. The protein was solvated in a rhombic dodecahedron with SPC water molecules, such that the minimal distance of the protein to the periodic boundary was 10 Å, and ions were added to neutralize the charge on the protein and simulate a 150 mM NaCl environment (68 Na+ ions and 62 Cl-ions). Energy minimization was achieved using the steepest descent algorithm. A 100 ps isothermal (NVT) MD simulation with position restraints on the protein was used to equilibrate the system at 300 K. A time step of 2 fs was used, and the system was equilibrated and simulated for a total of 100 ns with isotropic Parrinello-Rahman pressure coupling (τp = 12 ps, reference pressure = 1 bar) and V-rescale temperature coupling (300 K). Long-range electrostatics were calculated with the Particle Mesh Ewald (PME) method (cutoff = 1.2 nm) and van der Waals interactions employed a force-switch modifier with a cutoff of 1.2 nm. Following a 50 ns equilibration period, the four simulations were continued for 300 ns total.

### Structural analysis

Structural analysis was performed using LigPlot+ ^62,63^ and PHENIX to investigate protein-ligand interactions and overall structural quality. LigPlot+ was used to generate 2D diagrams of hydrogen bonds and hydrophobic interactions between the protein and ligand. The PHENIX suite was employed for model validation, including geometry analysis, clash detection, and refinement statistics to ensure structural accuracy.

### Flow cytometry

Ancestral constructs and reporter plasmids were transformed into *E. coli* 3.300 and plated onto LB supplemented with 100 mg/L each of ampicillin and chloramphenicol (Sigma). Three colonies of each plate were picked and used to inoculate 10 mL M9 media with 1% glycerol seed culture supplemented with 100 mg/L ampicillin and chloramphenicol. The seed cultures were incubated with shaking at 37 ºC for 24 hours. The OD_600_ of each culture was measured, and a 96-well plate prepared with each culture diluted to OD_600_ of 0.01 in minimal media with 1% glycerol to a total volume of 1 mL. Stock solutions of lactose (positive control), BMDG, D-fucose, or sucrose (negative control) were added to a final concentration of 7.5 mM per well. The plate layout was randomized, covered with breathable film, and incubated at 37 ºC for 18 hours. Cells were harvested by centrifuging (2000 x g for 10 minutes at 25 ºC) and resuspended in phosphate buffer (50 mM Na_2_HPO_4_, 150 mM NaCl, pH 7.4). Cell counting was conducted on a APF LSR II flow cytometer, and the results analyzed using FlowJo™ v10.8 Software (BD Life Sciences). Gating strategy shown in **Supplementary Figure 9**. Fold GFP induction for each sugar was calculated based on mean detected GFP fluorescence measured in triplicate, normalized to no ligand control.

## Acknowledgements

This work was supported by the Australian Research Council Centre of Excellence in Synthetic Biology (CE200100029) (CJJ & JAK), the Australian Research Council Centre of Excellence for Innovations in Protein and Peptide Science (CE200100012) (CJJ). This research was undertaken in part using the MX2 beamline at the Australian Synchrotron, part of ANSTO, and made use of the Australian Cancer Research Foundation (ACRF) detector. We acknowledge and thank the National Computing Infrastructure of Australia (NCI) for providing access to the Gadi supercomputer, and the cell sorting suite at the John Curtin School of Medical Research (JCSMR) for use of their flow cytometry facilities. RG was supported by a Deakin PhD Scholarship and a Rod Rickards PhD Scholarship.

## Author contributions

Conceptualization: RG, HB, MS, CJ,

Methodology: RG, HB, JK, RF, LT, NT, MS

Investigation: RG, HB, RF, LT, MS

Visualization: RG, MS

Funding acquisition: CJ

Project administration: CJ

Supervision: CJ

Writing – original draft: RG, HB, MS, CJ

Writing – review & editing: RG, JK, MS, CJ

## Additional information

Supplementary Information is available for this paper. Structures have been deposited to RSCB PDB under IDs 9NT3, 9NT4, 9NT5, 9NT6.

## References

1 Conant, G. C. & Wolfe, K. H. Turning a hobby into a job: How duplicated genes find new functions. Nature Reviews Genetics 9, 938–950 (2008). 10.1038/nrg2482

2 Khersonsky, O. & Tawfik, D. S. Enzyme promiscuity: a mechanistic and evolutionary perspective. Annu. Rev. Biochem. 79, 471–505 (2010). 10.1146/annurev-biochem-030409-143718

3 Näsvall, J., Sun, L., Roth, J. R. & Andersson, D. I. Real-Time Evolution of New Genes by Innovation, Amplification, and Divergence. Science 338, 384–387 (2012). 10.1126/science.1226521

4 Clifton, B. E. & Jackson, C. J. Ancestral Protein Reconstruction Yields Insights into Adaptive Evolution of Binding Specificity in Solute-Binding Proteins. Cell Chem Biol 23, 236–245 (2016). 10.1016/j.chembiol.2015.12.010

5 Glasner, M. E., Truong, D. P. & Morse, B. C. How enzyme promiscuity and horizontal gene transfer contribute to metabolic innovation. FEBS J 287, 1323–1342 (2020). 10.1111/febs.15185

6 Weickert, M. J. & Adhya, S. A family of bacterial regulators homologous to Gal and Lac repressors. J Biol Chem 267, 15869–15874 (1992).

7 Taraban, M. et al. Ligand-induced conformational changes and conformational dynamics in the solution structure of the lactose repressor protein. J Mol Biol 376, 466–481 (2008). 10.1016/j.jmb.2007.11.067

8 Glasgow, A. et al. Ligand-specific changes in conformational flexibility mediate long-range allostery in the lac repressor. Nature Communications 14, 1179 (2023). 10.1038/s41467-023-36798-1

9 Clifton, B. E. et al. Evolution of cyclohexadienyl dehydratase from an ancestral solute-binding protein. Nat. Chem. Biol. 14, 542–547 (2018). 10.1038/s41589-018-0043-2

10 Kaczmarski, J. A. et al. Altered conformational sampling along an evolutionary trajectory changes the catalytic activity of an enzyme. Nat. Commun. 11, 5945 (2020). 10.1038/s41467-020-19695-9

11 Fukami-Kobayashi, K., Tateno, Y. & Nishikawa, K. Parallel evolution of ligand specificity between LacI/GalR family repressors and periplasmic sugar-binding proteins. Mol. Biol. Evol. 20, 267–277 (2003). 10.1093/molbev/msg038

12 Taylor, N. D. et al. Engineering an allosteric transcription factor to respond to new ligands. Nat. Methods 13, 177–183 (2016). 10.1038/nmeth.3696

13 Wu, J. et al. Design and application of a lactulose biosensor. Sci. Rep. 7, 45994 (2017). 10.1038/srep45994

14 Juárez, J. F., Lecube-Azpeitia, B., Brown, S. L., Johnston, C. D. & Church, G. M. Biosensor libraries harness large classes of binding domains for construction of allosteric transcriptional regulators. Nat. Commun. 9, 3101 (2018). 10.1038/s41467-018-05525-6

15 Mannan, A. A., Liu, D., Zhang, F. & Oyarzún, D. A. Fundamental Design Principles for Transcription-Factor-Based Metabolite Biosensors. ACS Synth. Biol. 6, 1851–1859 (2017). 10.1021/acssynbio.7b00172

16 Kaczmarski, J. A., Mitchell, J. A., Spence, M. A., Vongsouthi, V. & Jackson, C. J. Structural and evolutionary approaches to the design and optimization of fluorescence-based small molecule biosensors. Curr. Opin. Struct. Biol. 57, 31–38 (2019). 10.1016/j.sbi.2019.01.013

17 Rondon, R. E., Groseclose, T. M., Short, A. E. & Wilson, C. J. Transcriptional programming using engineered systems of transcription factors and genetic architectures. Nature Communications 10, 4784 (2019). 10.1038/s41467-019-12706-4

18 Daber, R., Sochor, M. A. & Lewis, M. Thermodynamic Analysis of Mutant lac Repressors. Journal of Molecular Biology 409, 76–87 (2011). 10.1016/j.jmb.2011.03.057

19 Battistuzzi, F. U., Feijao, A. & Hedges, S. B. A genomic timescale of prokaryote evolution: insights into the origin of methanogenesis, phototrophy, and the colonization of land. BMC Evol Biol 4, 44 (2004). 10.1186/1471-2148-4-44

20 Meinhardt, S. et al. Novel insights from hybrid LacI/GalR proteins: family-wide functional attributes and biologically significant variation in transcription repression. Nucleic Acids Res 40, 11139–11154 (2012). 10.1093/nar/gks806

21 Swint-Kruse, L. & Matthews, K. S. Allostery in the LacI/GalR family: variations on a theme. Curr Opin Microbiol 12, 129–137 (2009). 10.1016/j.mib.2009.01.009

22 Ravcheev, D. A. et al. Comparative genomics and evolution of regulons of the LacI-family transcription factors. Front Microbiol 5, 294 (2014). 10.3389/fmicb.2014.00294

23 Meger, A. T. et al. Rugged fitness landscapes minimize promiscuity in the evolution of transcriptional repressors. Cell Syst 15, 374-387.e376 (2024). 10.1016/j.cels.2024.03.002

24 Spence, M. A., Kaczmarski, J. A., Saunders, J. W. & Jackson, C. J. Ancestral sequence reconstruction for protein engineers. Curr Opin Struct Biol 69, 131–141 (2021). 10.1016/j.sbi.2021.04.001

25 Le, S. Q. & Gascuel, O. An improved general amino acid replacement matrix. Mol Biol Evol 25, 1307–1320 (2008). 10.1093/molbev/msn067

26 Kalyaanamoorthy, S., Minh, B. Q., Wong, T. K. F., von Haeseler, A. & Jermiin, L. S. ModelFinder: fast model selection for accurate phylogenetic estimates. Nature Methods 14, 587–589 (2017). 10.1038/nmeth.4285

27 Shimodaira, H. An approximately unbiased test of phylogenetic tree selection. Syst. Biol. 51, 492–508 (2002). 10.1080/10635150290069913

28 Yang, Z. PAML 4: Phylogenetic Analysis by Maximum Likelihood. Molecular Biology and Evolution 24, 1586–1591 (2007). 10.1093/molbev/msm088

29 Pérez-Rueda, E. & Collado-Vides, J. The repertoire of DNA-binding transcriptional regulators in Escherichia coli K-12. Nucleic Acids Res 28, 1838–1847 (2000). 10.1093/nar/28.8.1838

30 Parente, D. J. & Swint-Kruse, L. Multiple co-evolutionary networks are supported by the common tertiary scaffold of the LacI/GalR proteins. PLoS One 8, e84398 (2013). 10.1371/journal.pone.0084398

31 Eick, G. N., Bridgham, J. T., Anderson, D. P., Harms, M. J. & Thornton, J. W. Robustness of Reconstructed Ancestral Protein Functions to Statistical Uncertainty. Mol Biol Evol 34, 247–261 (2017). 10.1093/molbev/msw223

32 McKellar, J. L. O., Minnell, J. J. & Gerth, M. L. A high-throughput screen for ligand binding reveals the specificities of three amino acid chemoreceptors from Pseudomonas syringae pv. actinidiae. Mol. Microbiol. 96, 694–707 (2015). 10.1111/mmi.12964

33 Williams, P. D., Pollock, D. D., Blackburne, B. P. & Goldstein, R. A. Assessing the accuracy of ancestral protein reconstruction methods. PLoS Comput. Biol. 2, e69 (2006). 10.1371/journal.pcbi.0020069

34 Tartèse, R., Chaussidon, M., Gurenko, A., Delarue, F. & Robert, F. Warm Archaean oceans reconstructed from oxygen isotope composition of early-life remnants. Geochem. Perspect. Lett., 55–65 (2017). 10.7185/geochemlet.1706

35 Sadler, J. R., Sasmor, H. & Betz, J. L. A perfectly symmetric lac operator binds the lac repressor very tightly. Proc. Natl. Acad. Sci. U. S. A. 80, 6785–6789 (1983). 10.1073/pnas.80.22.6785

36 Luria, S. E., Adams, J. N. & Ting, R. C. Transduction of lactose-utilizing ability among strains of E. coli and S. dysenteriae and the properties of the transducing phage particles. Virology 12, 348–390 (1960). 10.1016/0042-6822(60)90161-6

37 Chatterjee, S., Zhou, Y. N., Roy, S. & Adhya, S. Interaction of Gal repressor with inducer and operator: induction of gal transcription from repressor-bound DNA. Proc. Natl. Acad. Sci. U. S. A. 94, 2957–2962 (1997). 10.1073/pnas.94.7.2957

38 Cooper, G. M. The Cell: A Molecular Approach. (Sinauer Associates, 2000).

39 Donnér, J., Caruthers, M. H. & Gill, S. J. A calorimetric investigation of the interaction of the lac repressor with inducer. J Biol Chem 257, 14826–14829 (1982).

40 Wilson, C. J., Zhan, H., Swint-Kruse, L. & Matthews, K. S. Ligand interactions with lactose repressor protein and the repressor-operator complex: the effects of ionization and oligomerization on binding. Biophys Chem 126, 94–105 (2007). 10.1016/j.bpc.2006.06.005

41 Forman-Kay, J. D. The ‘dynamics’ in the thermodynamics of binding. Nat. Struct. Biol. 6, 1086–1087 (1999). 10.1038/70008

42 Wankowicz, S. & Fraser, J. Making sense of chaos: uncovering the mechanisms of conformational entropy. ChemRxiv (2024). 10.26434/chemrxiv-2023-9b5k7-v3

43 Caro, J. A. et al. Entropy in molecular recognition by proteins. Proc Natl Acad Sci U S A 114, 6563–6568 (2017). 10.1073/pnas.1621154114

44 Fukami-Kobayashi, K., Tateno, Y. & Nishikawa, K. Parallel Evolution of Ligand Specificity Between LacI/GalR Family Repressors and Periplasmic Sugar-Binding Proteins. Molecular Biology and Evolution 20, 267–277 (2003). 10.1093/molbev/msg038

45 Lin, J. & and Qian, T. Earth’s Climate History from 4.5 Billion Years to One Minute. Atmosphere-Ocean 60, 188–232 (2022). 10.1080/07055900.2022.2082914

46 Garcia, A. K., Schopf, J. W., Yokobori, S.-i., Akanuma, S. & Yamagishi, A. Reconstructed ancestral enzymes suggest long-term cooling of Earth’s photic zone since the Archean. Proceedings of the National Academy of Sciences 114, 4619–4624 (2017). 10.1073/pnas.1702729114

47 Gaucher, E. A., Govindarajan, S. & Ganesh, O. K. Palaeotemperature trend for Precambrian life inferred from resurrected proteins. Nature 451, 704–707 (2008). 10.1038/nature06510

48 Chang, C.-E. A., Chen, W. & Gilson, M. K. Ligand configurational entropy and protein binding. Proc Natl Acad Sci U S A 104, 1534–1539 (2007). 10.1073/pnas.0610494104

49 Aragão, D. et al. MX2: a high-flux undulator microfocus beamline serving both the chemical and macromolecular crystallography communities at the Australian Synchrotron. J Synchrotron Radiat 25, 885–891 (2018). 10.1107/s1600577518003120

50 Potterton, L. et al. CCP4i2: the new graphical user interface to the CCP4 program suite. Acta Crystallogr D Struct Biol 74, 68–84 (2018). 10.1107/S2059798317016035

51 Winn, M. D. et al. Overview of the CCP4 suite and current developments. Acta Crystallogr. D Biol. Crystallogr. 67, 235–242 (2011). 10.1107/S0907444910045749

52 McCoy, A. J. et al. Phaser crystallographic software. J Appl Crystallogr 40, 658–674 (2007). 10.1107/s0021889807021206

53 Jumper, J. et al. Highly accurate protein structure prediction with AlphaFold. Nature 596, 583–589 (2021). 10.1038/s41586-021-03819-2

54 Murshudov, G. N., Vagin, A. A. & Dodson, E. J. Refinement of macromolecular structures by the maximum-likelihood method. Acta Crystallogr D Biol Crystallogr 53, 240–255 (1997). 10.1107/s0907444996012255

55 Liebschner, D. et al. Macromolecular structure determination using X-rays, neutrons and electrons: recent developments in Phenix. Acta Crystallogr D Struct Biol 75, 861–877 (2019). 10.1107/s2059798319011471

56 Emsley, P., Lohkamp, B., Scott, W. G. & Cowtan, K. Features and development of Coot. Acta Crystallogr D Biol Crystallogr 66, 486–501 (2010). 10.1107/s0907444910007493

57 Croll, T. I. ISOLDE: a physically realistic environment for model building into low-resolution electron-density maps. Acta Crystallogr D Struct Biol 74, 519–530 (2018). 10.1107/S2059798318002425

58 Keller, S. et al. High-precision isothermal titration calorimetry with automated peak-shape analysis. Anal. Chem. 84, 5066–5073 (2012). 10.1021/ac3007522

59 Zhao, H., Piszczek, G. & Schuck, P. SEDPHAT--a platform for global ITC analysis and global multi-method analysis of molecular interactions. Methods 76, 137–148 (2015). 10.1016/j.ymeth.2014.11.012

60 Brautigam, C. A. Calculations and Publication-Quality Illustrations for Analytical Ultracentrifugation Data. Methods Enzymol. 562, 109–133 (2015). 10.1016/bs.mie.2015.05.001

61 Huang, J. et al. CHARMM36m: an improved force field for folded and intrinsically disordered proteins. Nat Methods 14, 71–73 (2017). 10.1038/nmeth.4067

62 Laskowski, R. A. & Swindells, M. B. LigPlot+: multiple ligand-protein interaction diagrams for drug discovery. J Chem Inf Model 51, 2778–2786 (2011). 10.1021/ci200227u

63 Wallace, A. C., Laskowski, R. A. & Thornton, J. M. LIGPLOT: a program to generate schematic diagrams of protein-ligand interactions. Protein Eng 8, 127–134 (1995). 10.1093/protein/8.2.127

