## Supplementary Information for "The temperature dependence of binding entropy is a selective pressure in protein evolution"

**Supplementary Figure 1.** Posterior probability distributions of Anc1-4. Each ancestral regulator was reconstructed with a high mean posterior probability (>0.8, averaged across all sites) and a small proportion of sites reconstructed with low posterior probability (<0.5).

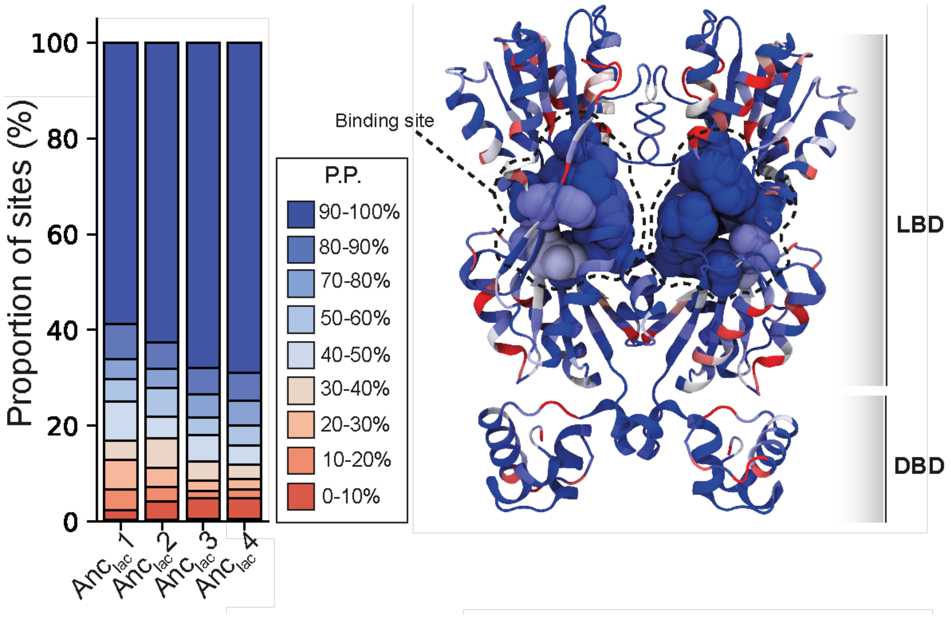

**Supplementary Figure 2.** Sequence alignment of ancestral TFs and ecLacI, aligned using MUSCLE [64] and visualized using ggmsa [65] in Rstudio.

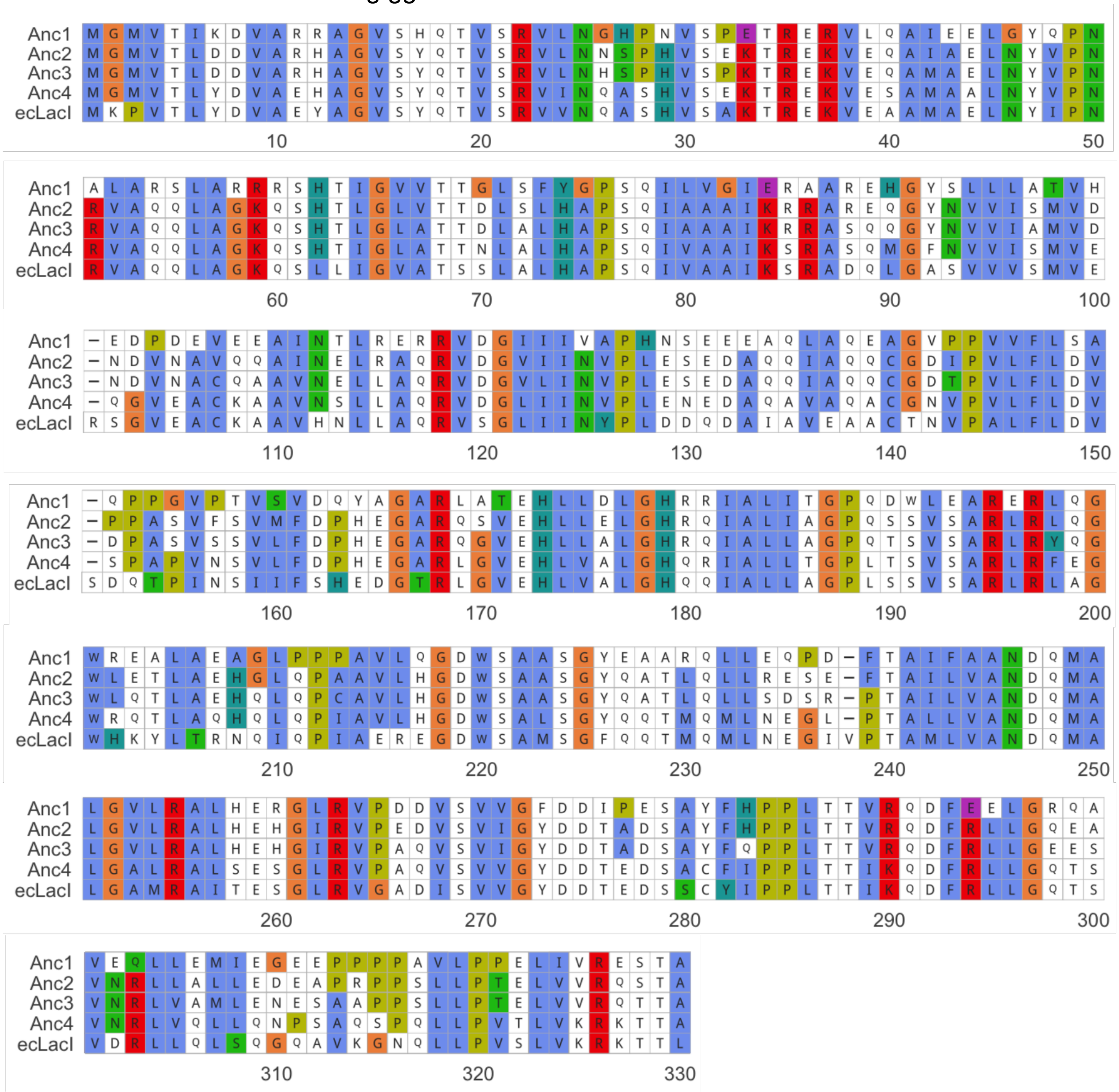

**Supplementary Figure 3.** Cartoon representation of mutations up the evolutionary trajectory, modelled as spheres on LacI (PDB ID 2P9H, shown in dark blue).

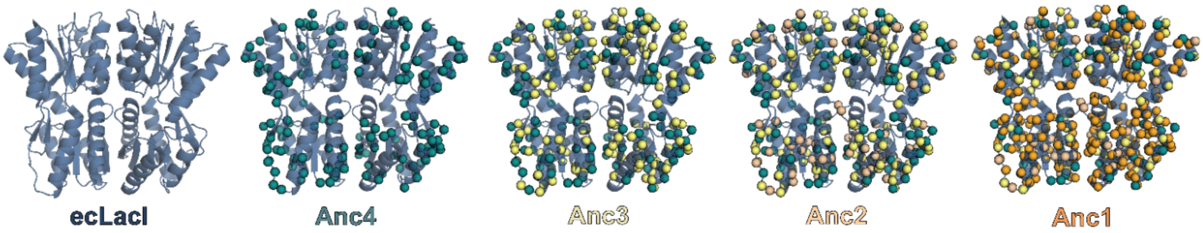

**Supplementary Figure 4.** Full configuration of Biolog Microbial Phenotyping plates PM1 and PM2A used for DSF screening of ancestral small molecule binding.

### PM1 Carbon Utilization Assays

Catalog #12111

| | | | | | | | | | | | |
|---|---|---|---|---|---|---|---|---|---|---|---|
| A1<br>Negative Control | A2<br>L-Arabinose | A3<br>N-Acetyl-D-Glucosamine | A4<br>D-Saccharic Acid | A5<br>Succinic Acid | A6<br>D-Galactose | A7<br>L-Aspartic Acid | A8<br>L-Proline | A9<br>D-Alanine | A10<br>D-Trehalose | A11<br>D-Mannose | A12<br>Dulcitol |
| B1<br>D-Serine | B2<br>D-Sorbitol | B3<br>Glycerol | B4<br>L-Fucose | B5<br>D-Glucuronic Acid | B6<br>D-Gluconic Acid | B7<br>D,L-α-Glycerol-Phosphate | B8<br>D-Xylose | B9<br>L-Lactic Acid | B10<br>Formic Acid | B11<br>D-Mannitol | B12<br>L-Glutamic Acid |
| C1<br>D-Glucose-6-Phosphate | C2<br>D-Galactonic Acid-γ-Lactone | C3<br>D,L-Malic Acid | C4<br>D-Ribose | C5<br>Tween 20 | C6<br>L-Rhamnose | C7<br>D-Fructose | C8<br>Acetic Acid | C9<br>α-D-Glucose | C10<br>Maltose | C11<br>D-Melibiose | C12<br>Thymidine |
| D-1<br>L-Asparagine | D2<br>D-Aspartic Acid | D3<br>D-Glucosaminic Acid | D4<br>1,2-Propanediol | D5<br>Tween 40 | D6<br>α-Keto-Glutaric Acid | D7<br>α-Keto-Butyric Acid | D8<br>α-Methyl-D-Galactoside | D9<br>α-D-Lactose | D10<br>Lactulose | D11<br>Sucrose | D12<br>Uridine |
| E1<br>L-Glutamine | E2<br>m-Tartaric Acid | E3<br>D-Glucose-1-Phosphate | E4<br>D-Fructose-6-Phosphate | E5<br>Tween 80 | E6<br>α-Hydroxy Glutaric Acid-γ-Lactone | E7<br>α-Hydroxy Butyric Acid | E8<br>ß-Methyl-D-Glucoside | E9<br>Adonitol | E10<br>Maltotriose | E11<br>2-Deoxy Adenosine | E12<br>Adenosine |
| F1<br>Glycyl-L-Aspartic Acid | F2<br>Citric Acid | F3<br>myo-Inositol | F4<br>D-Threonine | F5<br>Fumaric Acid | F6<br>Bromo Succinic Acid | F7<br>Propionic Acid | F8<br>Mucic Acid | F9<br>Glycolic Acid | F10<br>Glyoxylic Acid | F11<br>D-Cellobiose | F12<br>Inosine |
| G1<br>Glycyl-L-Glutamic Acid | G2<br>Tricarballylic Acid | G3<br>L-Serine | G4<br>L-Threonine | G5<br>L-Alanine | G6<br>L-Alanyl-Glycine | G7<br>Acetoacetic Acid | G8<br>N-Acetyl-ß-D-Mannosamine | G9<br>Mono Methyl Succinate | G10<br>Methyl Pyruvate | G11<br>D-Malic Acid | G12<br>L-Malic Acid |
| H1<br>Glycyl-L-Proline | H2<br>p-Hydroxy Phenyl Acetic Acid | H3<br>m-Hydroxy Phenyl Acetic Acid | H4<br>Tyramine | H5<br>D-Psicose | H6<br>L-Lyxose | H7<br>Glucuronamide | H8<br>Pyruvic Acid | H9<br>L-Galactonic Acid-γ-Lactone | H10<br>D-Galacturonic Acid | H11<br>Phenylethyl-amine | H12<br>2-Aminoethanol |

### PM2A Carbon Utilization Assays

Catalog #12112

| | | | | | | | | | | | |
|---|---|---|---|---|---|---|---|---|---|---|---|
| A1<br>Negative Control | A2<br>Chondroitin Sulfate C | A3<br>α-Cyclodextrin | A4<br>ß-Cyclodextrin | A5<br>γ-Cyclodextrin | A6<br>Dextrin | A7<br>Gelatin | A8<br>Glycogen | A9<br>Inulin | A10<br>Laminarin | A11<br>Mannan | A12<br>Pectin |
| B1<br>N-Acetyl-D-Galactosamine | B2<br>N-Acetyl-Neuraminic Acid | B3<br>ß-D-Allose | B4<br>Amygdalin | B5<br>D-Arabinose | B6<br>D-Arabitol | B7<br>L-Arabitol | B8<br>Arbutin | B9<br>2-Deoxy-D-Ribose | B10<br>i-Erythritol | B11<br>D-Fucose | B12<br>3-0-ß-D-Galactopyranosyl-D-Arabinose |
| C1<br>Gentiobiose | C2<br>L-Glucose | C3<br>Lactitol | C4<br>D-Melezitose | C5<br>Maltitol | C6<br>α-Methyl-D-Glucoside | C7<br>ß-Methyl-D-Galactoside | C8<br>3-Methyl Glucose | C9<br>ß-Methyl-D-Glucuronic Acid | C10<br>α-Methyl-D-Mannoside | C11<br>ß-Methyl-D-Xyloside | C12<br>Palatinose |
| D1<br>D-Raffinose | D2<br>Salicin | D3<br>Sedoheptulosan | D4<br>L-Sorbose | D5<br>Stachyose | D6<br>D-Tagatose | D7<br>Turanose | D8<br>Xylitol | D9<br>N-Acetyl-D-Glucosaminitol | D10<br>γ-Amino Butyric Acid | D11<br>δ-Amino Valeric Acid | D12<br>Butyric Acid |
| E1<br>Capric Acid | E2<br>Caproic Acid | E3<br>Citraconic Acid | E4<br>Citramalic Acid | E5<br>D-Glucosamine | E6<br>2-Hydroxy Benzoic Acid | E7<br>4-Hydroxy Benzoic Acid | E8<br>ß-Hydroxy Butyric Acid | E9<br>Glycolic Acid | E10<br>α-Keto-Valeric Acid | E11<br>Itaconic Acid | E12<br>5-Keto-D-Gluconic Acid |
| F1<br>D-Lactic Acid Methyl Ester | F2<br>Malonic Acid | F3<br>Melibionic Acid | F4<br>Oxalic Acid | F5<br>Oxalomalic Acid | F6<br>Quinic Acid | F7<br>D-Ribono-1,4-Lactone | F8<br>Sebacic Acid | F9<br>Sorbic Acid | F10<br>Succinamic Acid | F11<br>D-Tartaric Acid | F12<br>L-Tartaric Acid |
| G1<br>Acetamide | G2<br>L-Alaninamide | G3<br>N-Acetyl-L-Glutamic Acid | G4<br>L-Arginine | G5<br>Glycine | G6<br>L-Histidine | G7<br>L-Homoserine | G8<br>Hydroxy-LProline | G9<br>L-Isoleucine | G10<br>L-Leucine | G11<br>L-Lysine | G12<br>L-Methionine |
| H1<br>L-Ornithine | H2<br>L-Phenylalanine | H3<br>L-Pyroglutamic Acid | H4<br>L-Valine | H5<br>D,L-Carnitine | H6<br>Sec-Butylamine | H7<br>D,L-Octopamine | H8<br>Putrescine | H9<br>Dihydroxy Acetone | H10<br>2,3-Butanediol | H11<br>2,3-Butanedione | H12<br>3-Hydroxy-2-Butanone |

**Supplementary Figure 5.** DSF screens of Anc1-4 and LacI against a library of small molecules. Error bars represent the standard error measurement from three replicates. **a.** Anc1 **b.** Anc2 **c.** Anc3 **d.** Anc4 **e.** ecLacI

**a.**

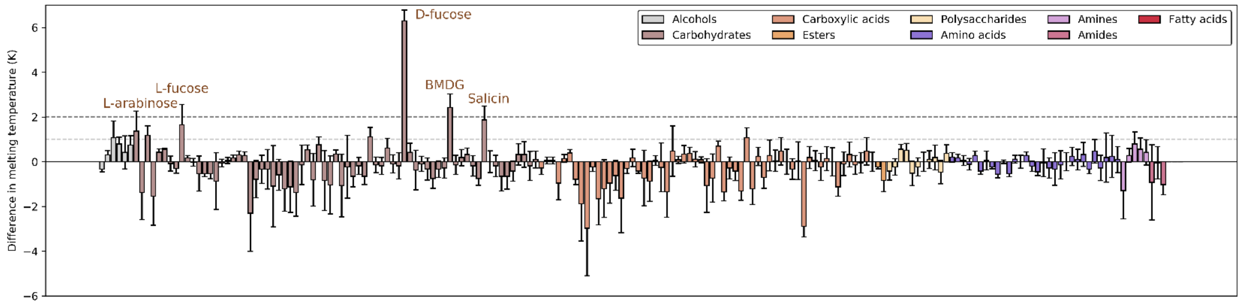


**b.**

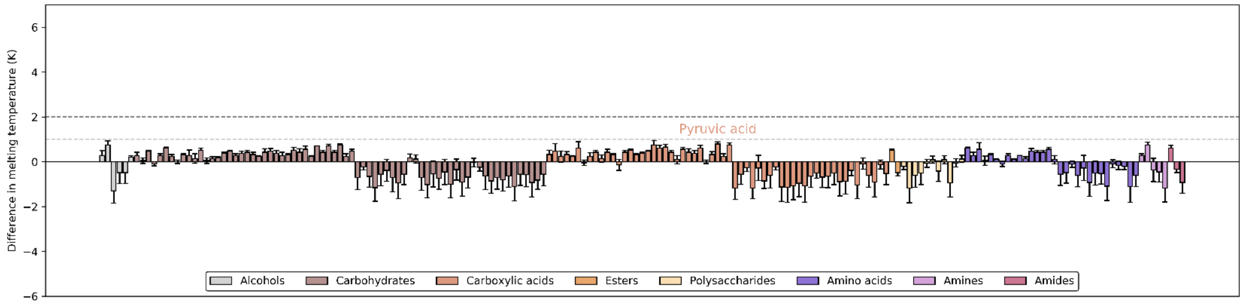


**c.**

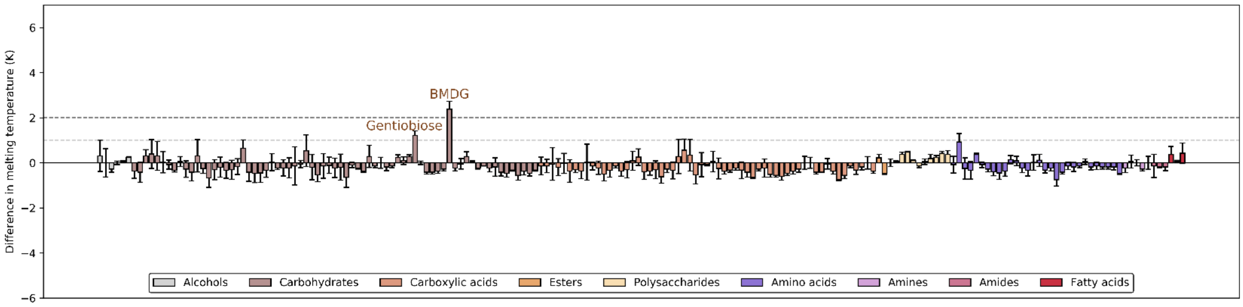


**d.**

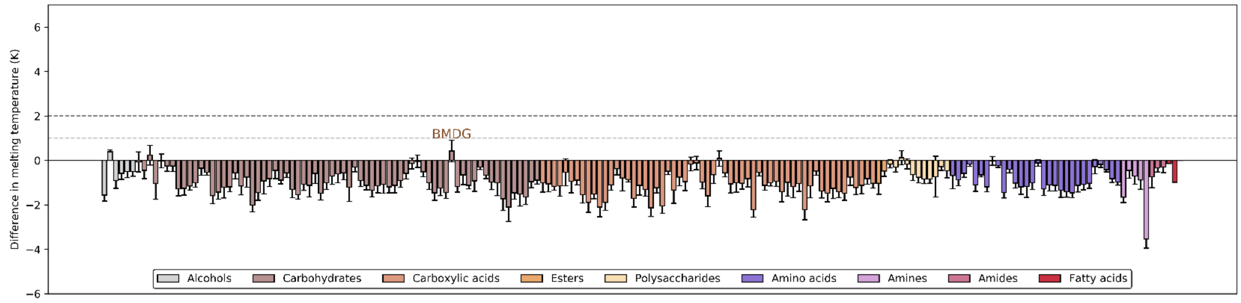


**e.**

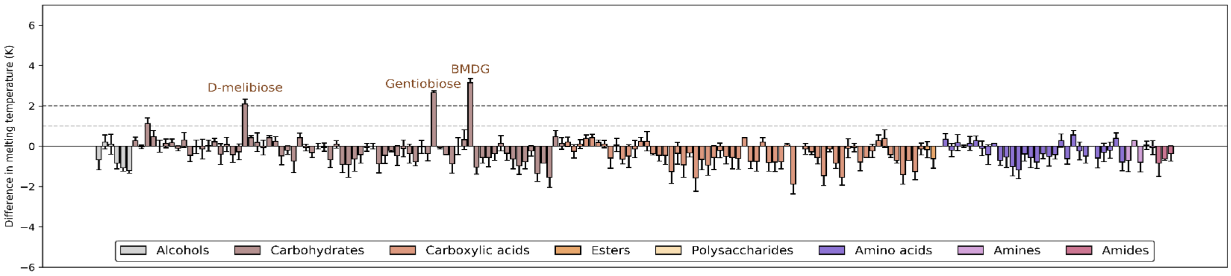

Difference in melting temperature (K)
6
4
2
0
−2
−4
−6
D-melibiose
Gentiobiose
BMDG
Alcohols
Carbohydrates
Carboxylic acids
Esters
Polysaccharides
Amino acids
Amines
Amides

**Supplementary Figure 6.** CD melting curves of ancestral transcription factors and calculated melting temperatures ($T_m$).

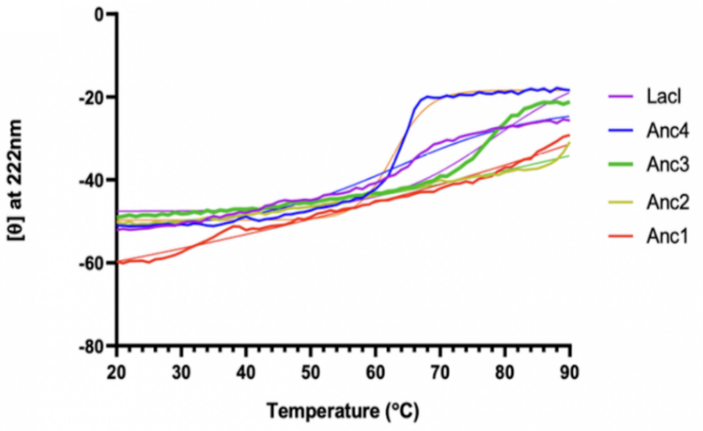


| Protein | $T_m$ CD ± SEM (ºC) | $T_m$ DSF ± SEM (ºC) |
|---|---|---|
| **Anc1** | 90.98 ± 0.45 | >100 |
| **Anc2** | 80.85 ± 6.03 | 87.4 ± 0.3 |
| **Anc3** | 78.04 ± 2.92 | 71.4 ± 0.4 |
| **Anc4** | 65.58 ± 0.70 | 56.9 ± 0.1 |
| **LacI** | 65.60 ± 1.53 | 58.8 ± 0.1 |

**Supplementary Figure 7.** Schematic of dual-plasmid reporter system and detailed plasmid maps.

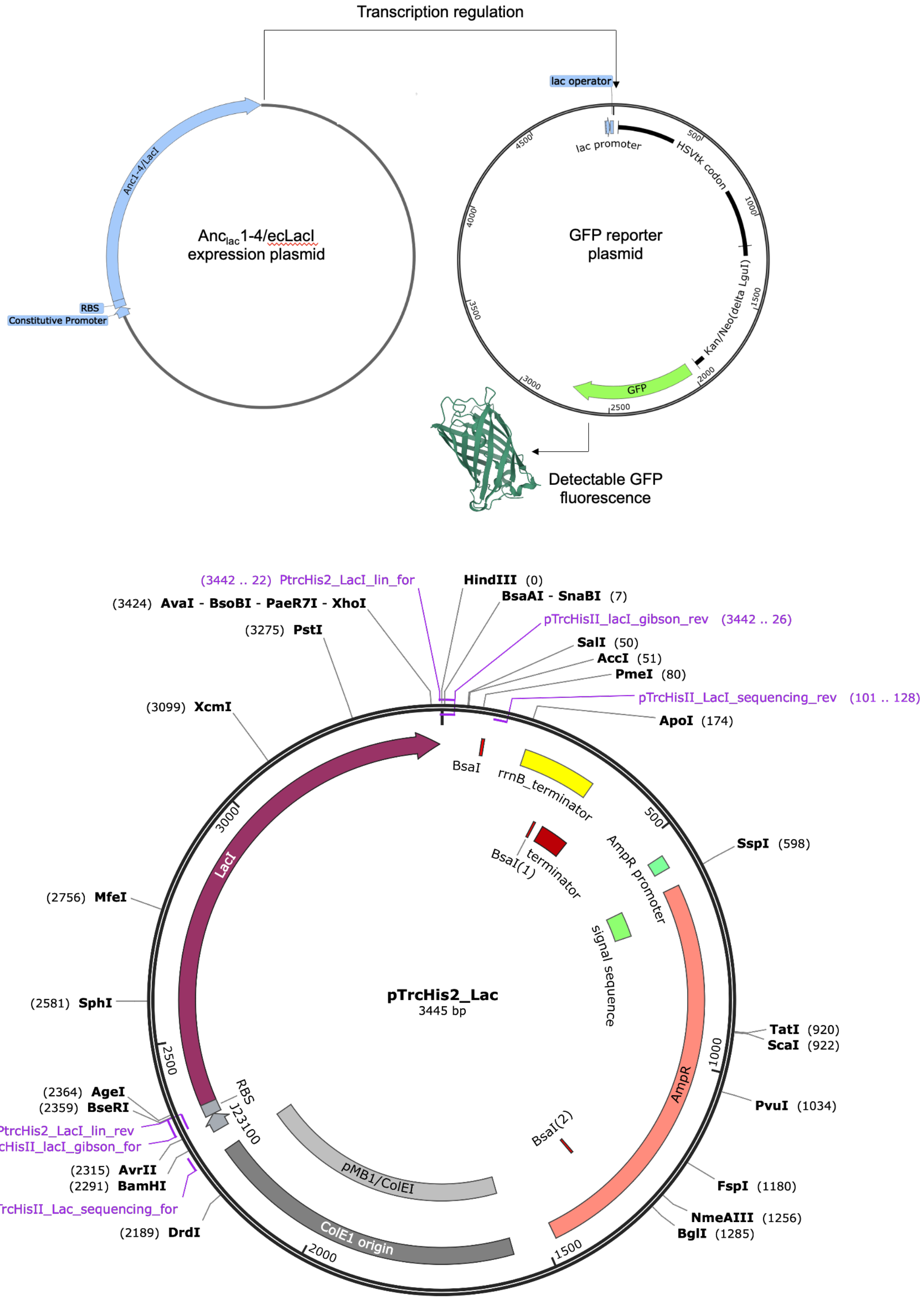

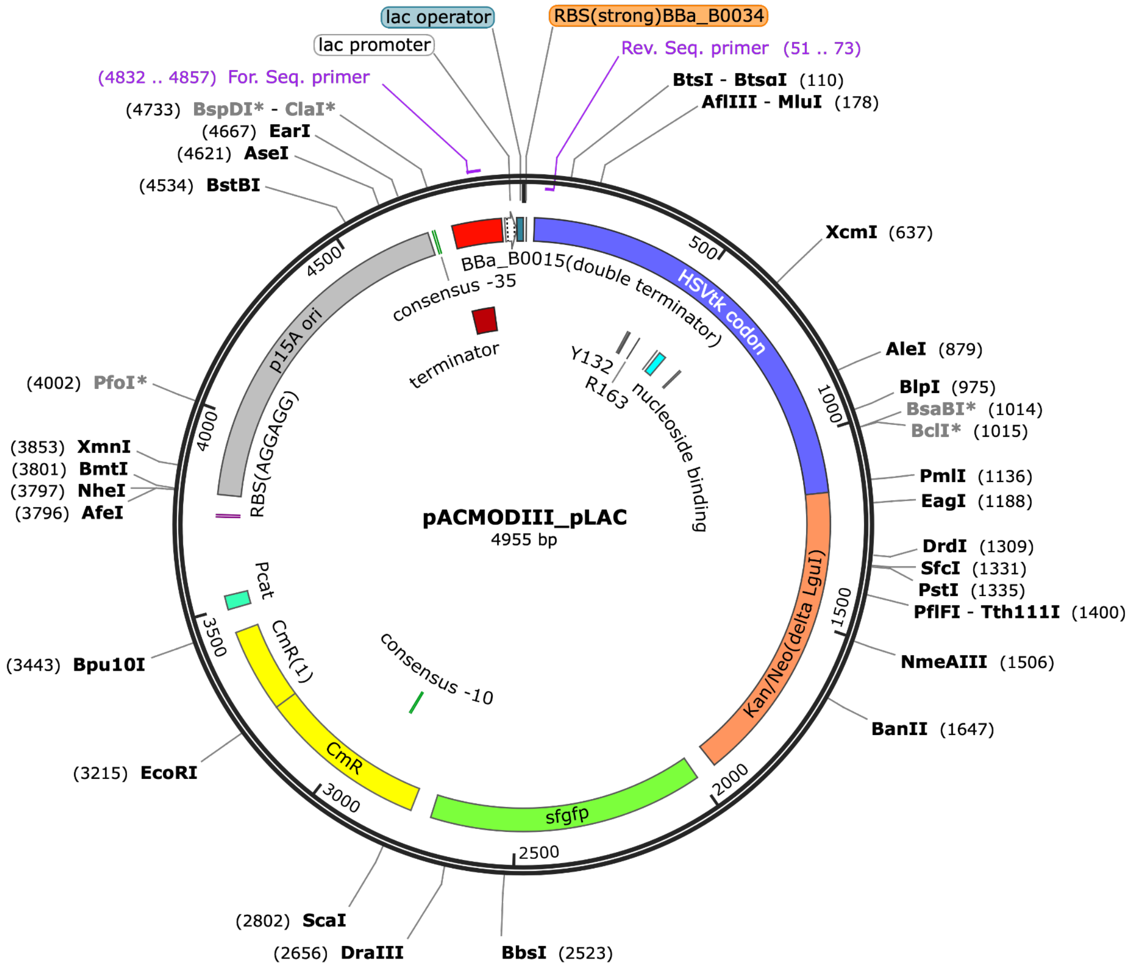

lac operator
RBS(strong)BBa_B0034
lac promoter
Rev. Seq. primer (51 .. 73)
(4832 .. 4857) For. Seq. primer
BtsI - BtsαI (110)
AflIII - MluI (178)
(4733) BspDI* - ClaI*
(4667) EarI
(4621) AseI
(4534) BstBI
XcmI (637)
AleI (879)
BlpI (975)
BsaBI* (1014)
BclI* (1015)
PmlI (1136)
EagI (1188)
DrdI (1309)
SfcI (1331)
PstI (1335)
PflFI - Tth111I (1400)
NmeAIII (1506)
BanII (1647)
BbsI (2523)
(2656) DraIII
(2802) ScaI
(3215) EcoRI
(3443) Bpu10I
(3796) AfeI
(3797) NheI
(3801) BmtI
(3853) XmnI
(4002) PfoI*
BBa_B0015(double terminator)
consensus -35
terminator
HSVtk codon
Y132
R163
nucleoside binding
p15A ori
RBS(AGGAGG)
pACMODIII_pLAC
4955 bp
Kan/Neo(delta LguI)
Pcat
CmR(1)
CmR
consensus -10
sfgfp
500
1000
1500
2000
2500
3000
3500
4000
4500

**Supplementary Figure 8.** Fold GFP induction, shown as the mean of three replicates (except Anc1 lactose and sucrose only two usable replicates due to low cell growth in minimal media) with SEM for each ancestor and tested sugar including positive and negative controls. Reported data is normalized to no ligand control, where sucrose and no ligand give almost identical responses.

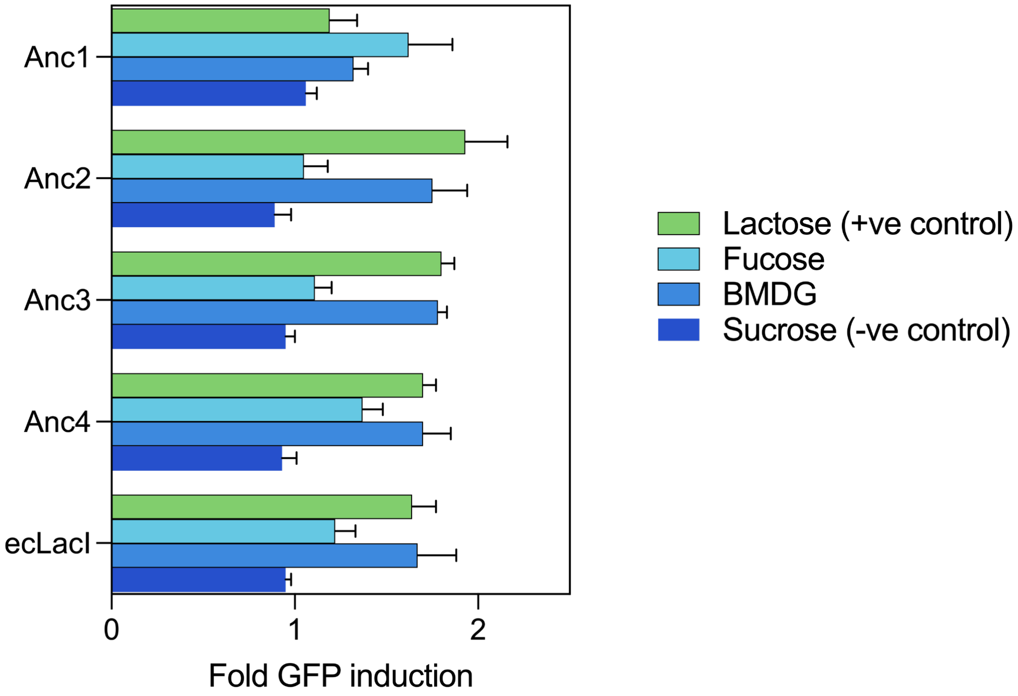

**Supplementary Figure 9. Gating strategies for flow cytometry.** Shown on the PACMODIII plasmid only (100% fluorescence, GFP not repressed). The mean Alexa Fluor 488-A (GFP) was used for calculation of GFP-fold induction by comparison of ligand-added to negative controls (sucrose only and no ligand).

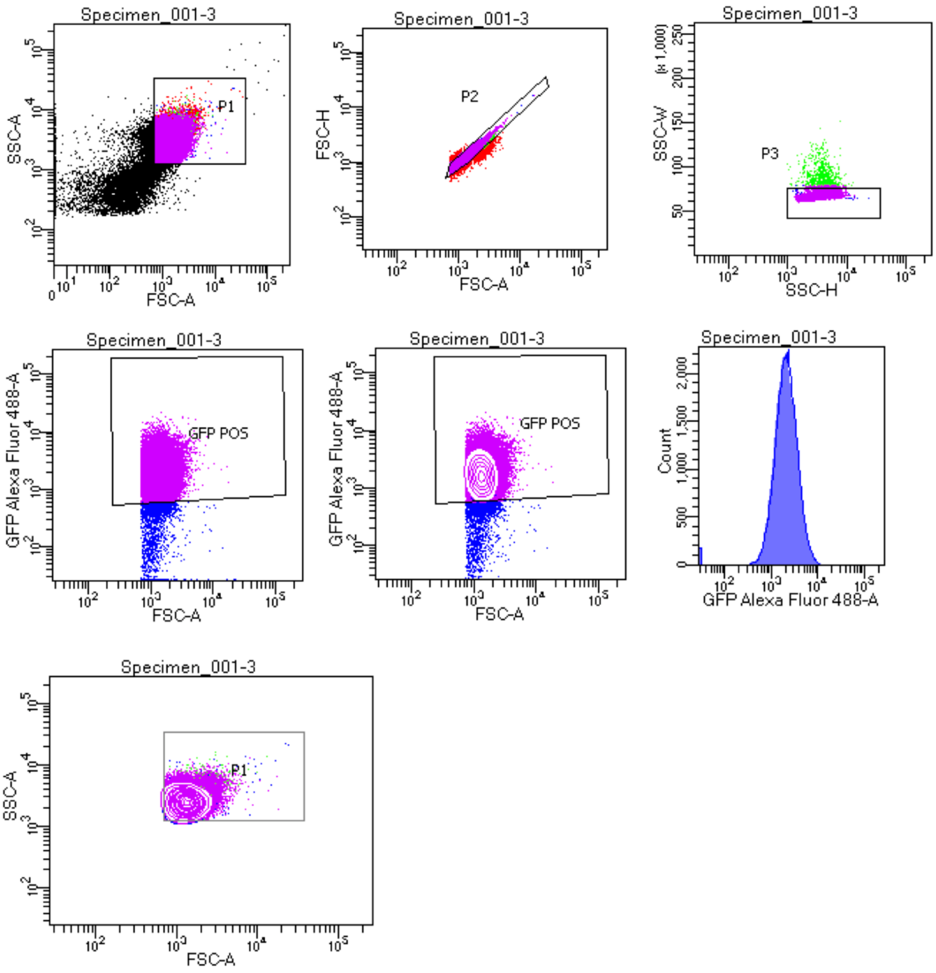


Tube: 3

| Population | #Events | %Parent | %Total |
|---|---|---|---|
| All Events | 100,000 | #### | 100.0 |
| P1 | 88,004 | 88.0 | 88.0 |
| P2 | 83,565 | 95.0 | 83.6 |
| P3 | 82,821 | 99.1 | 82.8 |
| GFP POS | 80,078 | 96.7 | 80.1 |

**Supplementary Figure 10. Flow cytometry histograms for each ancestor.** Using the above gating strategy for each biological replicate, the fold-GFP induction and chi-squared values were used to indicate statistically significant induction by ligands acting on the ancestral transcription factors. Representative histograms are shown.

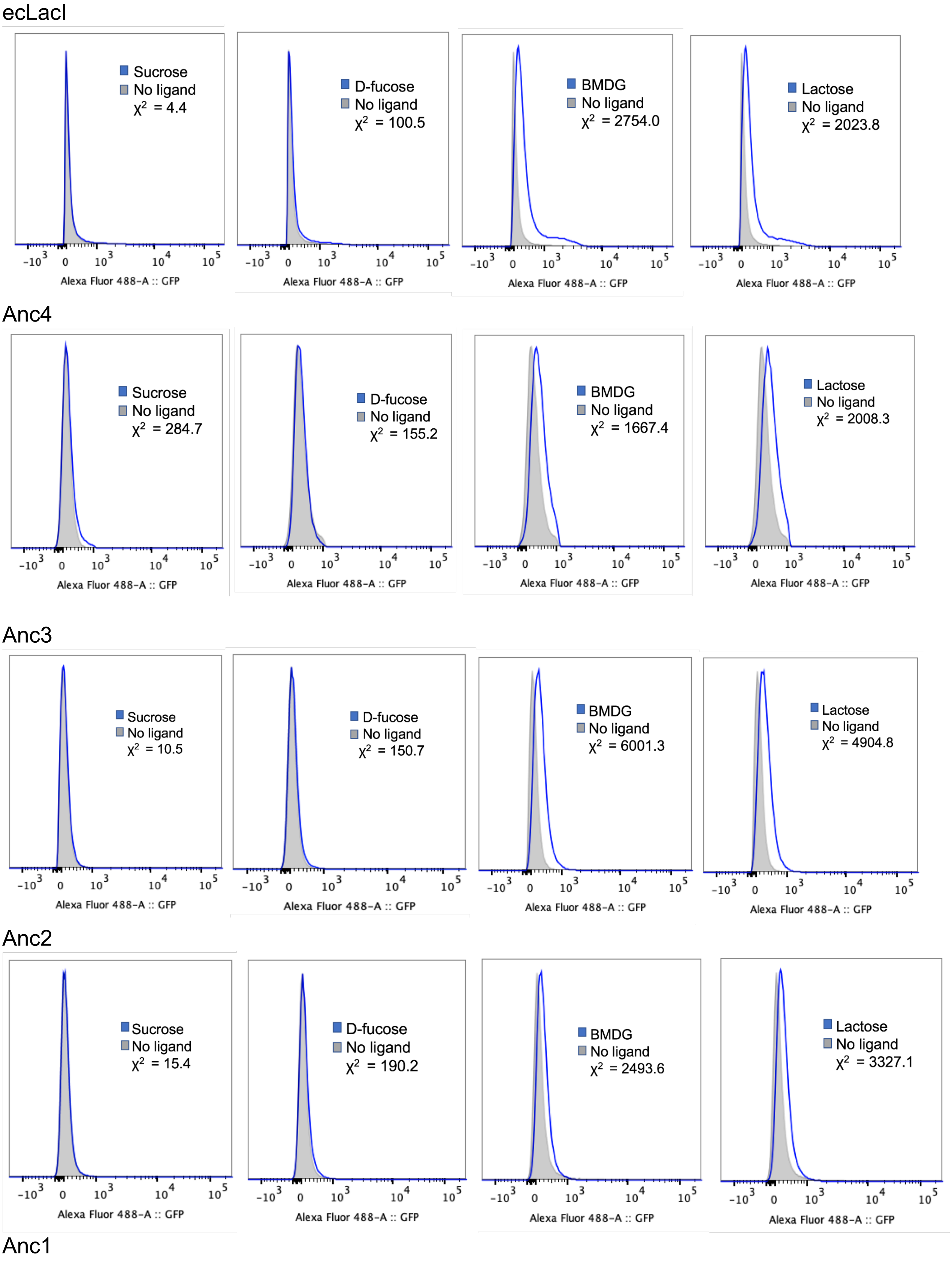

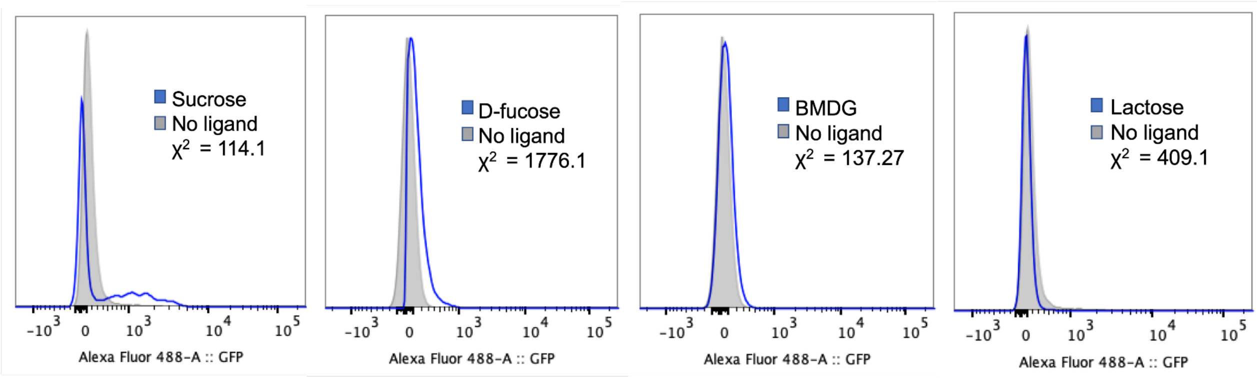
Sucrose
No ligand
χ2 = 114.1
D-fucose
No ligand
χ2 = 1776.1
BMDG
No ligand
χ2 = 137.27
Lactose
No ligand
χ2 = 409.1
-10^3 0 10^3 10^4 10^5
Alexa Fluor 488-A :: GFP

**Supplementary Table 1.** Full ITC data for ancestors.

| **Protein** | **Ligand** | **$K_D$ (mM)** | **$\Delta$H kJ/mol)** | **T$\Delta$S (kJ/mol)** | **$\Delta$G (kJ/mol)** | **Number of technical replicates** |
|---|---|---|---|---|---|---|
| **Anc1** | BMDG | 4.221 (3.608, 4.928) | 8.53 (6.70, 11.15) | 22.08 | -13.6 | 3 |
| **Anc2** | BMDG | 2.190 (1.847, 2.69) | 3.01 (NLF, 3.87) | 18.18 | -15.18 | 5 |
| **Anc3** | BMDG | 2.535 (2.325, 2.771) | -5.29 (-4.94, -5.67) | 9.53 | -14.82 | 4 |
| **Anc4** | BMDG | 0.550 (0.486, 0.619) | -4.63 (-4.15, -5.22) | 13.98 | -18.61 | 4 |
| **ecLacI** | BMDG | 0.306 (0.273, 0.367) | -24.9 (-22.85, <-83) | -4.87 | -20.05 | 3 |
| **Anc1** | D-fucose | 0.127 (0.110, 0.147) | 8.54 (7.25, 10.30) | 30.79 | -22.25 | 7 |
| **Anc4** | D-fucose | 14.74 (9.934, 19.840) | -6.91 (-4.68, -15.72) | 1.91 | -10.44 | 6 |
| **ecLacI** | D-fucose | 36.611 (24.605, 51.823) | -33.88 (<-50.2, >-20.9) | -25.68 | -8.2 | 5 |

Results from ITC analysis of Anc1-4 and LacI titrated BMDG and Anc1, Anc4 and LacI with D-fucose. Values in brackets are the lower and upper 68.3% confidence intervals (approximately equivalent to 1 S.D.) obtained from global analysis of technical repeats at 25 ºC. NLP = no limit found.

**Supplementary Figure 11.**

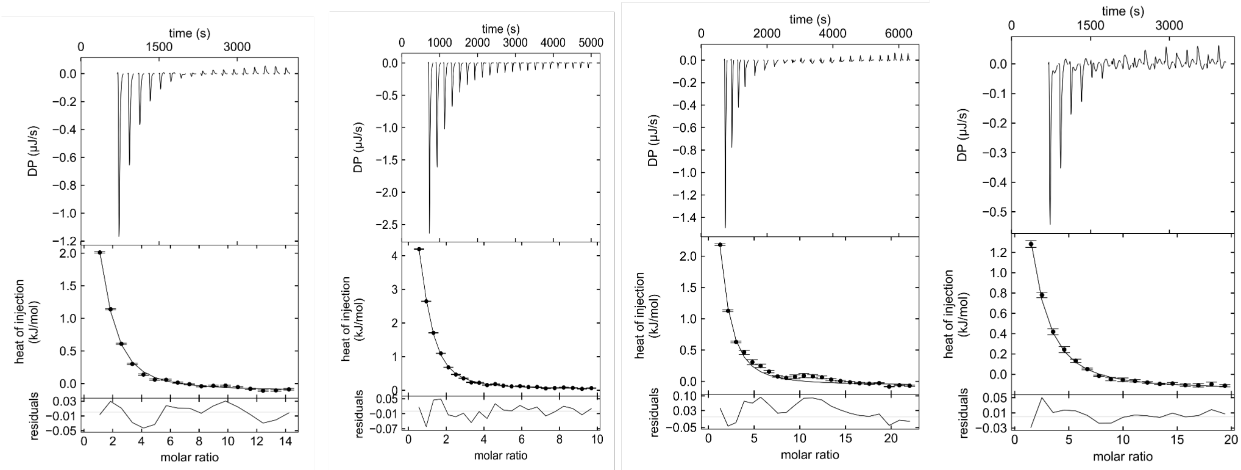


**ITC thermograms of titration of ᴅ-fucose into Anc1. (L-R). 1.** Cell = 151 µM Anc1 (monomer concentration); syringe = 8.9 mM ᴅ-fucose. **2.** Cell = 332 µM Anc1 (monomer concentration); syringe = 10.3 mM ᴅ-fucose. **3.** Cell =146 µM Anc1 (monomer concentration); syringe = 10.3 mM ᴅ-fucose. **4.** Cell = 83 µM Anc1 (monomer concentration); syringe = 6.7 mM ᴅ-fucose. The upper panels represent baseline-corrected power traces. The middle panels represent the integrated heat data and best fit of the independent binding site model in SEDPHAT (*48*). Figures were produced using GUSSI (*49*). The bottom panels show the residuals of the fit. Error bars represent the standard error in the integration of the peaks as calculated by NITPIC (*47*, *49*).

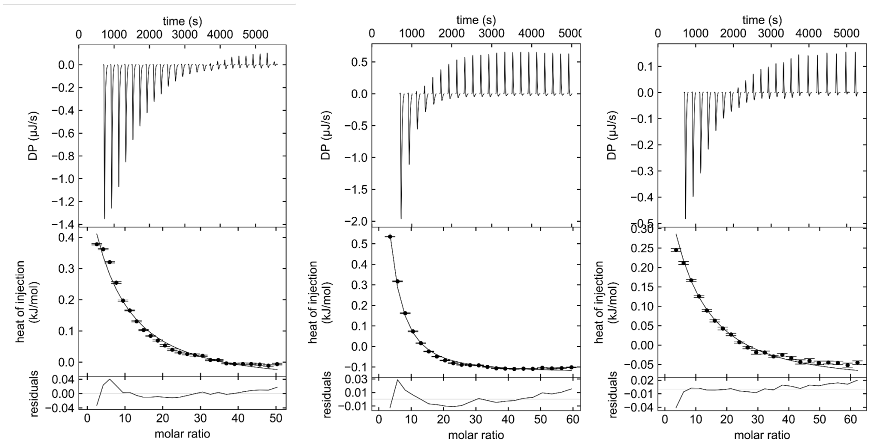


**ITC thermograms of titration of BMDG into Anc1. (L-R). 1.** Cell = 384 µM Anc1 (monomer concentration); syringe = 53.2 mM BMDG. **2.** Cell = 315 µM Anc1 (monomer concentration); syringe = 58.3 mM BMDG. **3.** Cell = 151 µM Anc1 (monomer concentration); syringe = 29.2 mM BMDG.

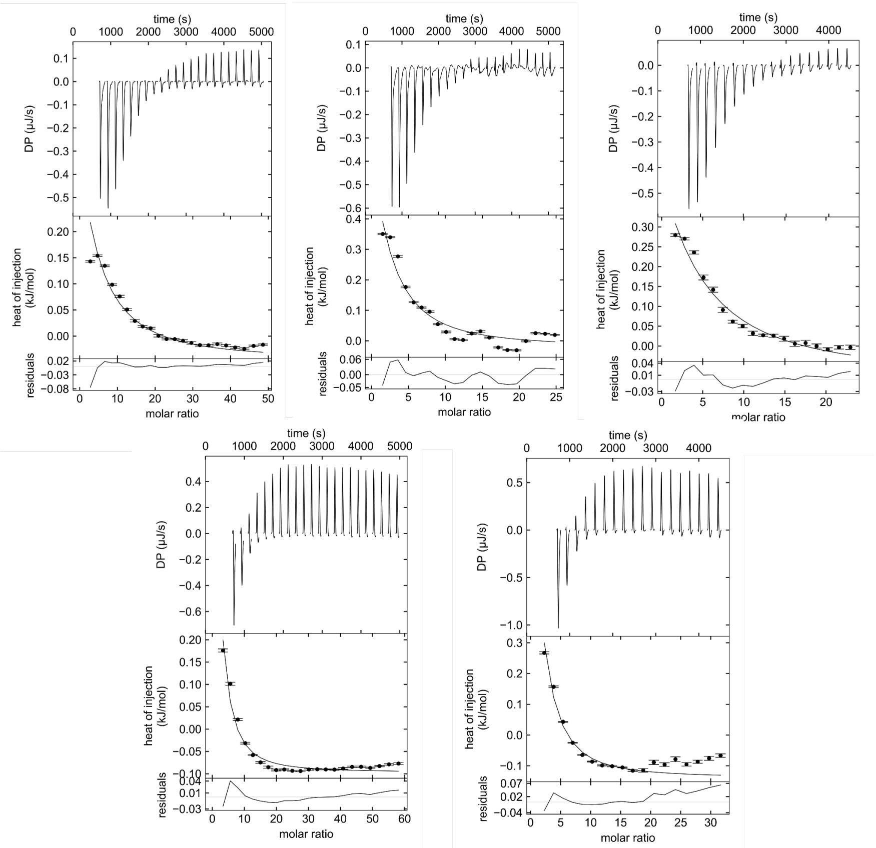


**ITC thermograms of titration of BMDG into Anc2. (L-R). 1.** Cell = 352 µM Anc2 (monomer concentration); syringe = 53.2 mM BMDG. 2. Cell = 359 µM Anc2 (monomer concentration); syringe = 29.2 mM BMDG. 3. Cell = 327 µM Anc2 (monomer concentration); syringe = 29.2 mM BMDG. 4. Cell = 323 µM Anc2 (monomer concentration); syringe = 58.3 mM BMDG. 5. Cell = 474 µM Anc2 (monomer concentration); syringe = 58.3 mM BMDG.

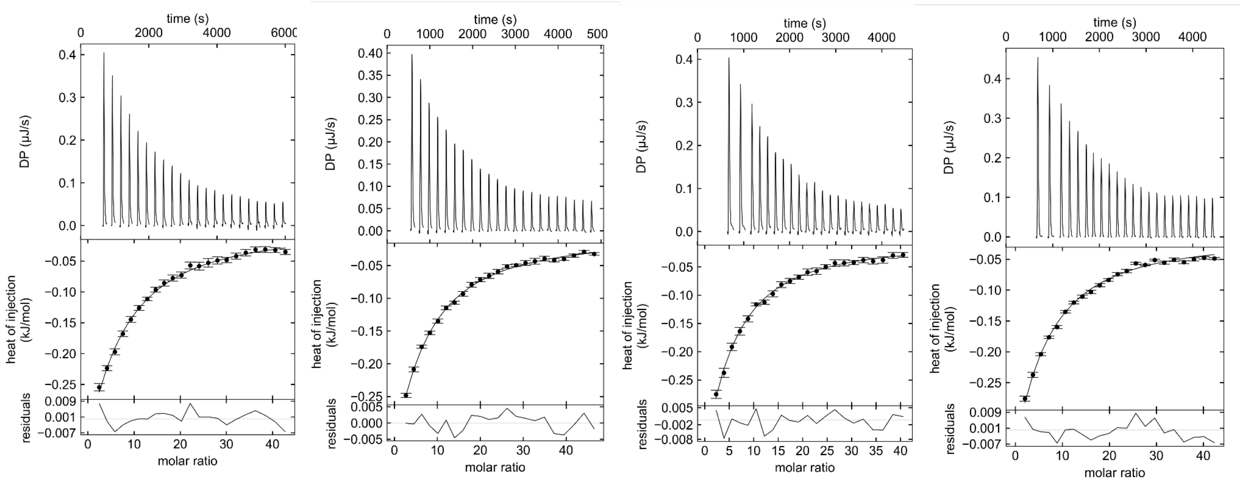


**ITC thermograms of titration of BMDG into Anc3. (L-R). 1.** Cell = 161 µM Anc3 (monomer concentration); syringe = 21.4 mM BMDG. **2.** Cell = 146 µM Anc3 (monomer concentration); syringe = 21.4 mM BMDG. **3.** Cell = 162 µM Anc3 (monomer concentration); syringe = 20.4 mM BMDG. **4.** Cell = 153 µM Anc3 (monomer concentration); syringe = 20.4 mM BMDG.

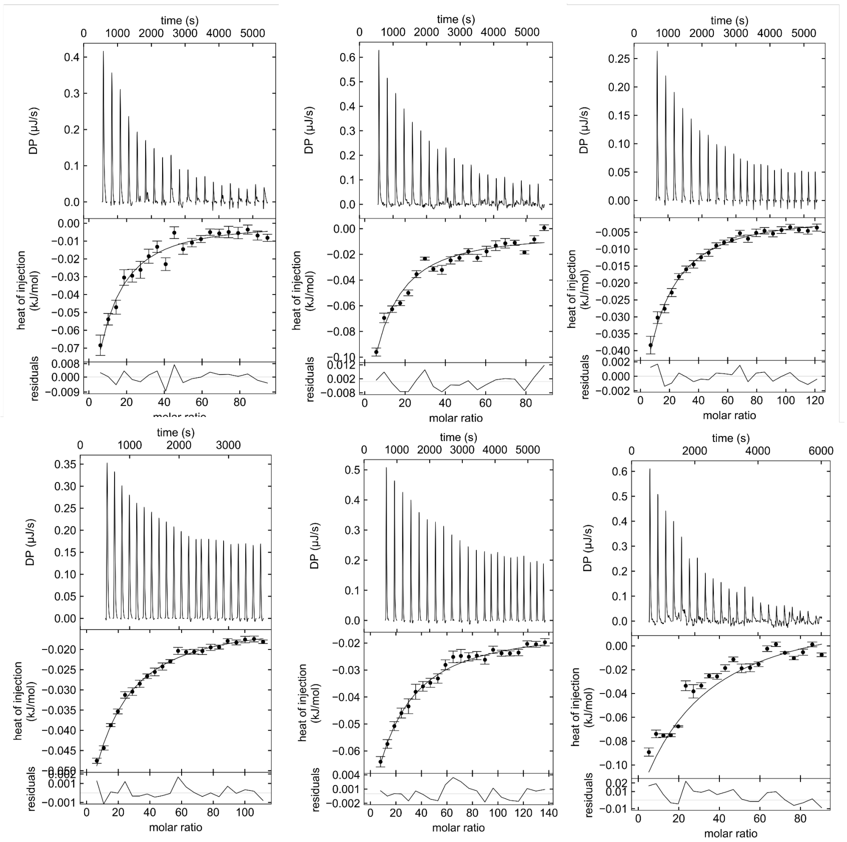


**ITC thermograms of titration of ᴅ-fucose into Anc4. (L-R). 1.** Cell = 273 µM Anc4 (monomer concentration); syringe = 89.4 mM ᴅ-fucose. **2.** Cell = 288 µM Anc4 (monomer concentration); syringe = 89.4 mM ᴅ-fucose. **3.** Cell = 237 µM Anc4 (monomer concentration); syringe = 89.4 mM ᴅ-fucose. **4.** Cell = 258 µM Anc4 (monomer concentration); syringe = 89.4 mM ᴅ-fucose. **5.** Cell = 210 µM Anc4 (monomer concentration); syringe = 89.4 mM ᴅ-fucose. **6.** Cell = 318 µM Anc4 (monomer concentration); syringe = 89.4 mM ᴅ-fucose.

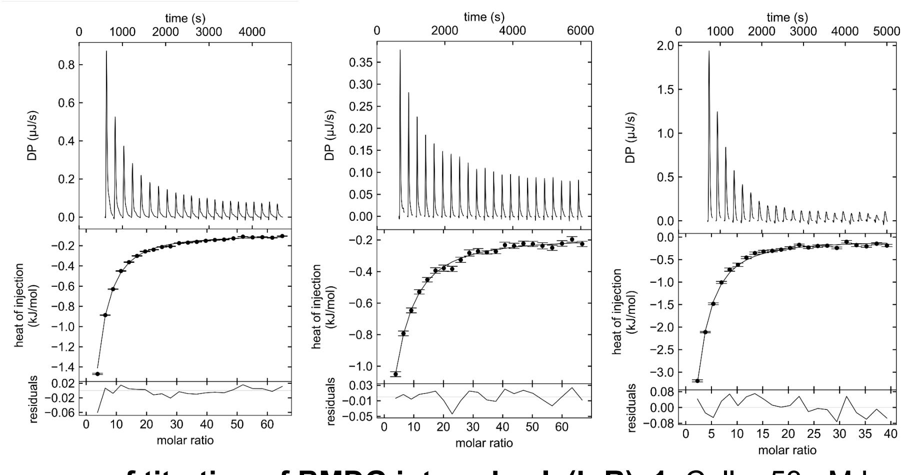


**ITC thermograms of titration of BMDG into ecLacI. (L-R). 1.** Cell = 53 µM LacI (monomer concentration); syringe = 107 mM BMDG. 2. Cell = 26 µM LacI (monomer concentration); syringe = 54.0 mM BMDG. 3. Cell = 82 µM LacI (monomer concentration); syringe = 100 mM BMDG.

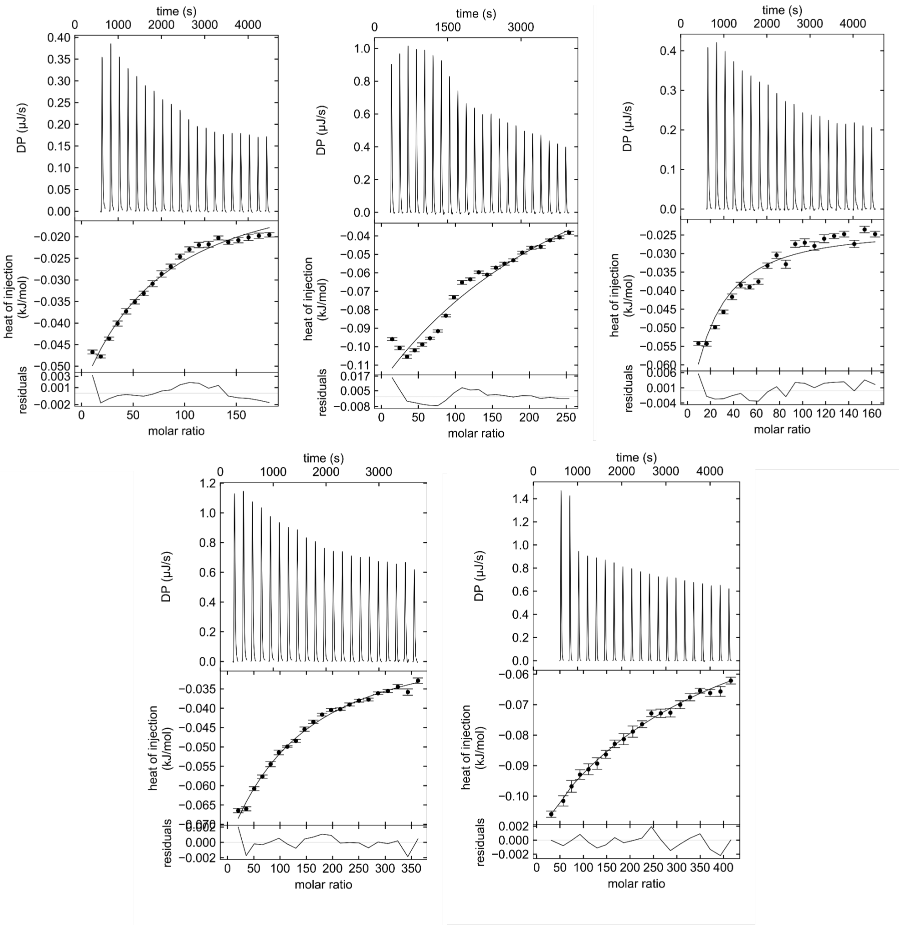


**ITC thermograms of titration of D-fucose into ecLacI. (L-R). 1.** Cell = 154 μM LacI (monomer concentration); syringe = 97.8 mM D-fucose. **2.** Cell = 128 μM LacI (monomer concentration); syringe = 101 mM D-fucose. **3.** Cell = 172 μM LacI (monomer concentration); syringe = 97.8 mM D-fucose. **4.** Cell = 169 μM LacI (monomer concentration); syringe = 202 mM D-fucose. **5.** Cell = 71 μM LacI (monomer concentration); syringe = 97.8 mM D-fucose.

**Supplementary Table 2.** Structure data collection and refinement statistics

| | **Anc1 apo** | **Anc1 D-fucose** | **Anc1 glycerol** | **Anc4 BMDG** |
|---|---|---|---|---|
| **PDB ID** | 9NT3 | 9NT4 | 9NT5 | 9NT6 |
| **Data collection** | | | | |
| Space group | P $2_1$ $2_1$ $2_1$ | P 1 $2_1$ 1 | P $3_1$ 2 1 | P $2_1$ $2_1$ $2_1$ |
| Cell dimensions | | | | |
| *a, b, c (Å)* | 62.86 85.59 211.78 | 63.45 211.0 87.32 | 104.82 104.82 50.13 | 36.47 108.34 119.60 |
| α, β, γ (°) | 90.0 90.0 90.0 | 90.0 90.51 90.0 | 90.0 90.0 120.0 | 90.0 90.0 90.0 |
| Resolution *(Å)* | 42.79-2.45 (2.54-2.45) | 43.66-2.40 (2.44-2.40) | 45.39-1.67 (1.70-1.67) | 49.34-1.50 (1.53-1.50) |
| $R_{merge}$ | 0.349 (2.515) | 0.199 (1.285) | 0.062 (5.031) | 0.140 (1.767) |
| $R_{pim}$ | 0.099 (0.697) | 0.080 (0.509) | 0.014 (1.122) | 0.041 (0.552) |
| I/σI | 6.8 (1.3) | 8.4 (1.6) | 22.5 (0.7) | 10.8 (1.4) |
| $CC_{1/2}$ | 0.994 (0.361) | 0.994 (0.395) | 1.000 (0.362) | 0.998 (0.653) |
| Completeness (%) | 100.0 (100.0) | 99.1 (98.6) | 100.0 (100.0) | 99.9 (98.3) |
| Redundancy | 13.4 (13.9) | 7.1 (7.3) | 20.6 (21.0) | 12.8 (11.0) |
| **Refinement** | | | | |
| Resolution *(Å)* | 42.79-2.45 (2.51-2.45) | 43.66-2.40 (2.43-2.40) | 34.31-1.67 (1.72-1.67) | 49.34-1.50 (1.52-1.50) |
| No. reflections | 42900 (4233) | 88448 (8773) | 37009 (3678) | 76787 (7495) |
| $R_{work}/R_{free}$ | 0.2243/0.2736 (0.3362/0.4021) | 0.2187/0.2701 (0.2933/0.3247) | 0.1887/0.2121 (0.3309/0.3369) | 0.1756/0.2078 (0.2415/0.2804) |
| No. atoms | 8184 | 16153 | 2244 | 4746 |
| Protein | 8105 | 15998 | 2098 | 4269 |
| Ligand/ion | 0 | 88 | 6 | 68 |
| Water | 79 | 67 | 140 | 409 |
| B-factors (overall) | 52.56 | 53.02 | 78.40 | 26.55 |
| Protein | 52.68 | 53.19 | 79.91 | 26.41 |
| Ligand/ion | - | 36.72 | 42.00 | 22.75 |
| Water | 40.43 | 32.70 | 57.33 | 28.67 |
| R.m.s. deviations | | | | |
| Bond lengths *(Å)* | 0.005 | 0.005 | 0.004 | 0.007 |
| Bond angles (°) | 0.743 | 0.688 | 0.719 | 0.988 |

**Supplementary Table 3.** ITC data for truncated ancestors

| Protein | Ligand | $K_D$ (mM) | $\Delta$H (kJ/mol) | $\Delta$S (J/mol/K) | $\Delta$G (kJ/mol) | Number of technical replicates |
|---|---|---|---|---|---|---|
| ΔAnc1 | D-fucose | 0.42 (0.32, 0.55) | 15.4 (13.6, 17.6) | 116.4 | -19.3 | 2 |
| ΔAnc4 | BMDG | 0.43 (0.34, 0.55) | -2.6 (-2.8, -2.4) | 55.6 | -19.2 | 2 |

Results from ITC analysis of truncated Δancestors show comparable thermodynamic binding parameters to full-length ancestors. Values in brackets are the lower and upper 68.3% confidence intervals (approx. 1 S.D.) obtained from global analysis of two technical replicates.

**Supplementary Figure 12.**

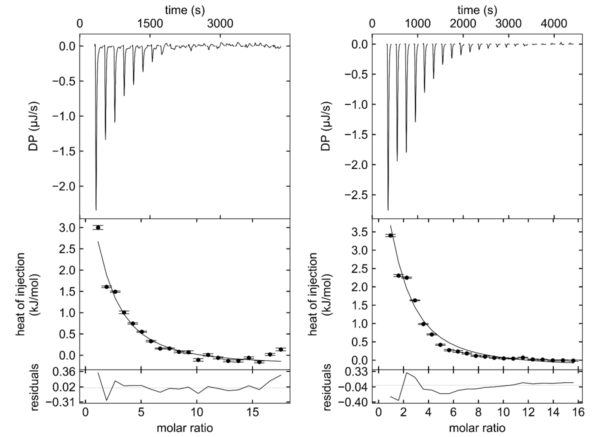


**ITC thermograms of titration of D-fucose into ΔAnc1. (L-R). 1.** Cell = 169 µM Anc1 (monomer concentration); syringe = 10 mM D-fucose. **2.** Cell = 141 µM Anc1 (monomer concentration); syringe = 11 mM D-fucose.

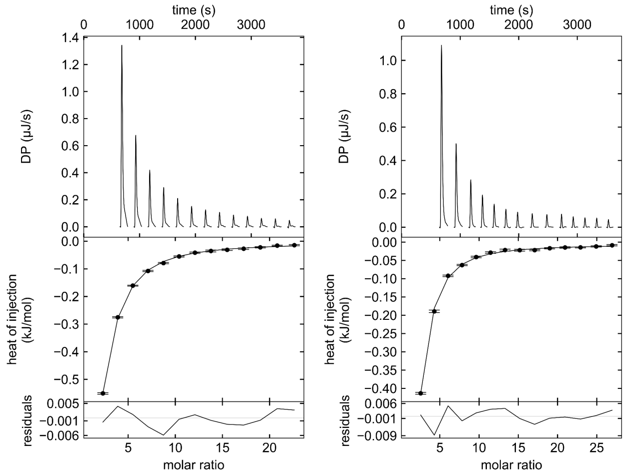


**ITC thermograms of titration of BMDG into ΔAnc4. (L-R). 1.** Cell = 275 µM Anc4 (monomer concentration); syringe = 38 mM BMDG. **2.** Cell = 303 µM Anc4 (monomer concentration); syringe = 38 mM BMDG.

**Supplementary Figure 13.** Circular dichroism (CD) and differential scanning fluorimetry (DSF) show the truncated ancestors (Anc1-4 ligand binding domain, LBD) are folded and form similar secondary structures to the full-length ancestral TFs.

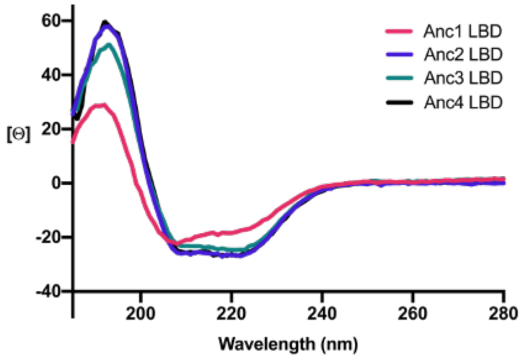


CD far-UV spectra obtained of Anc1–Anc4 LBDs at 20 ºC shows secondary structure consistent with α-helices and β-sheets.

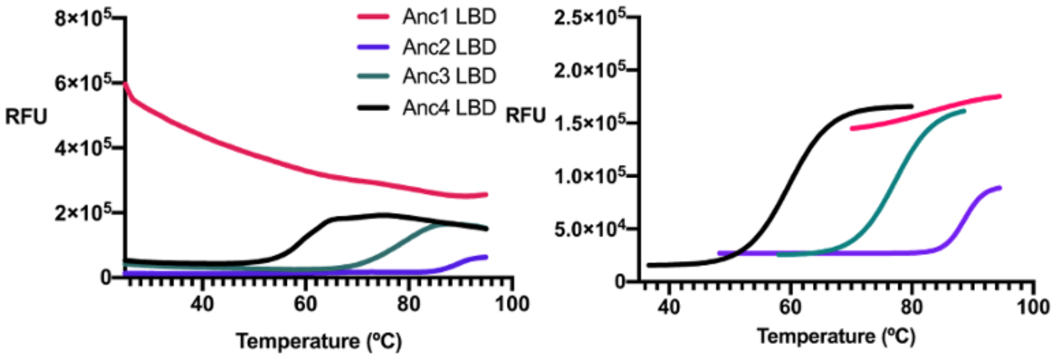


DSF melt curves of Anc1–Anc4 LBDs, RFU = relative fluorescence units. **Left.** Raw melt curves show dye binding in native state to Anc1 LBD. **Right.** Curve plotted using the Boltzmann model gives the melting temperature of Anc2, Anc3, Anc4 LBDs at the midpoint of the inflection.

**Supplementary Figure 14.** Binding pocket of LacI (blue) bound to IPTG showing residues.

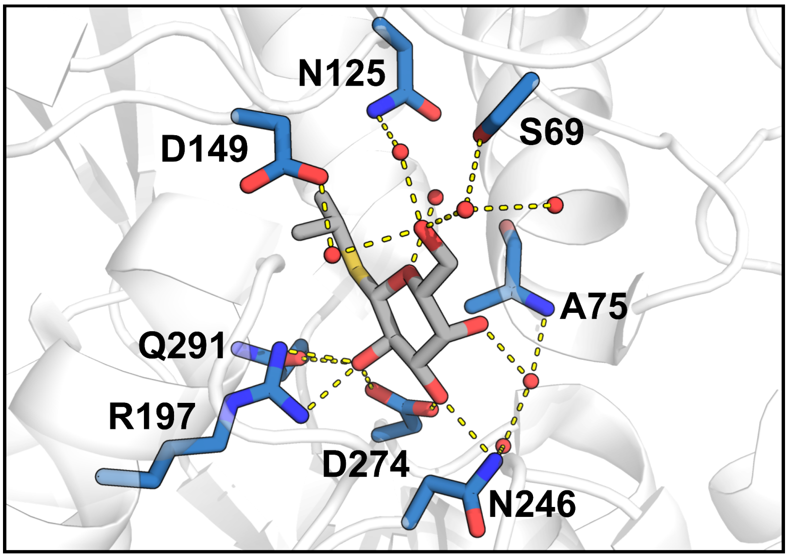

**Supplementary Figure 15.** RMSD of MD simulations of Anc4 LBD *apo.*

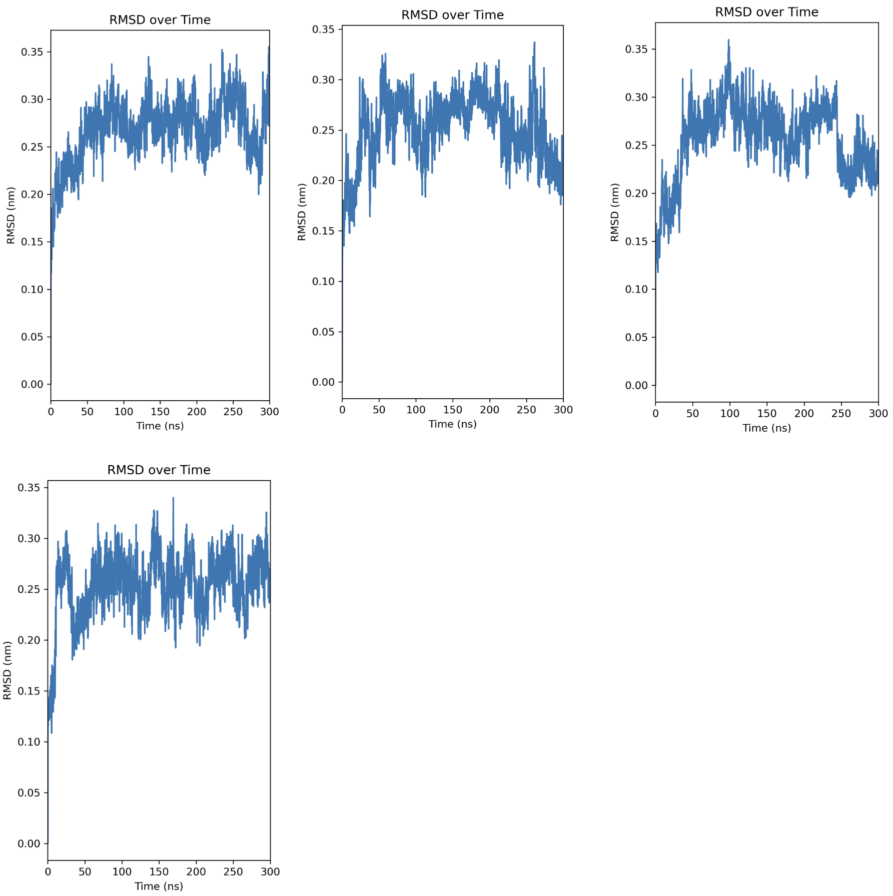

**Supplementary Figure 16.** ΔAnc1:ᴅ-fucose full crystal unit, of 4 dimeric pairs.

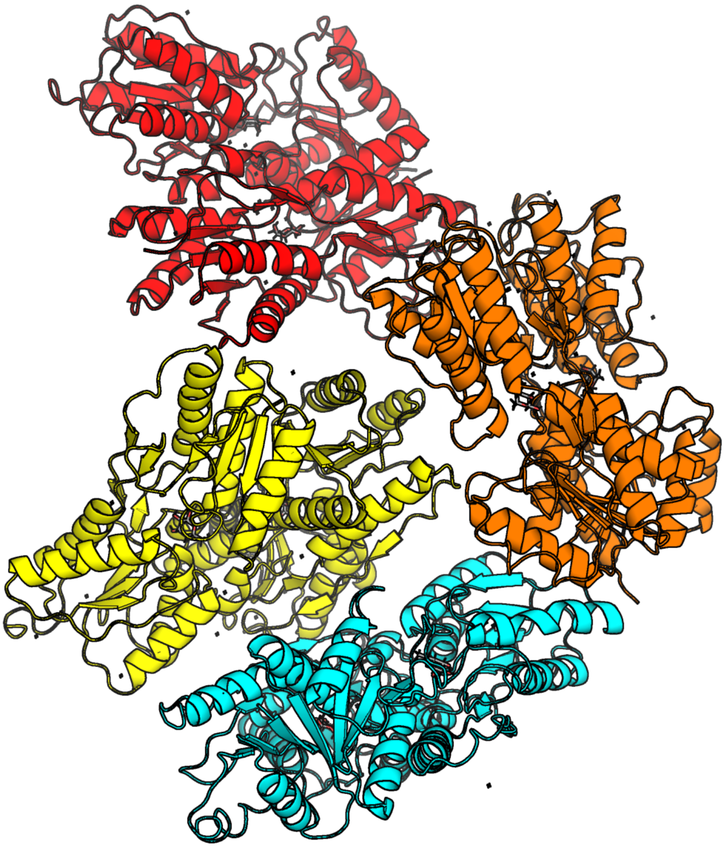

**Supplementary Figure 17.** LigPlot diagram of Anc1 and D-fucose, showing hydrogen bonds and hydrophobic interactions.

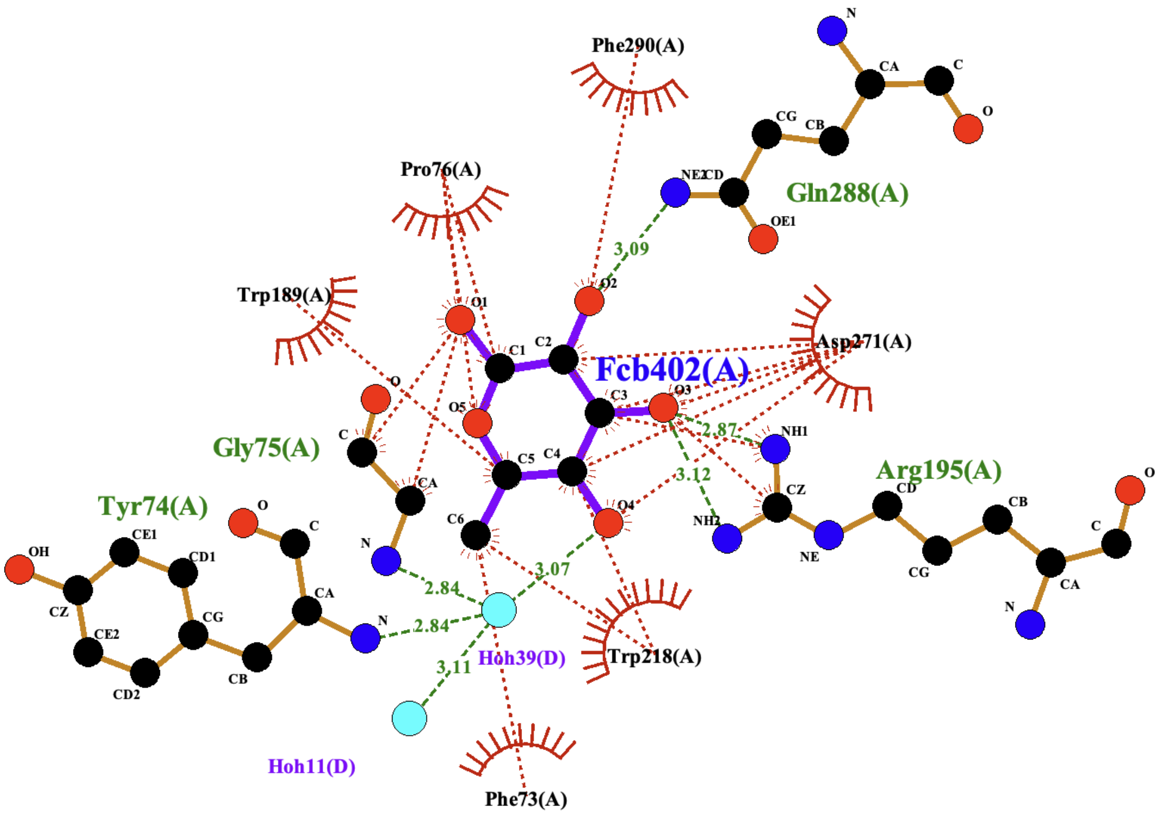

**Supplementary Figure 18.** Intrinsic tryptophan fluorescence of Anc1-Y74H mutant. **a.** Anc1:BMDG binding curve **b.** Anc1-Y74H mutant binding BMDG, no detectable curve **c.** Anc1:ᴅ-fucose binding curve **d.** Anc1-Y74H mutant binding ᴅ-fucose, no detectable curve.

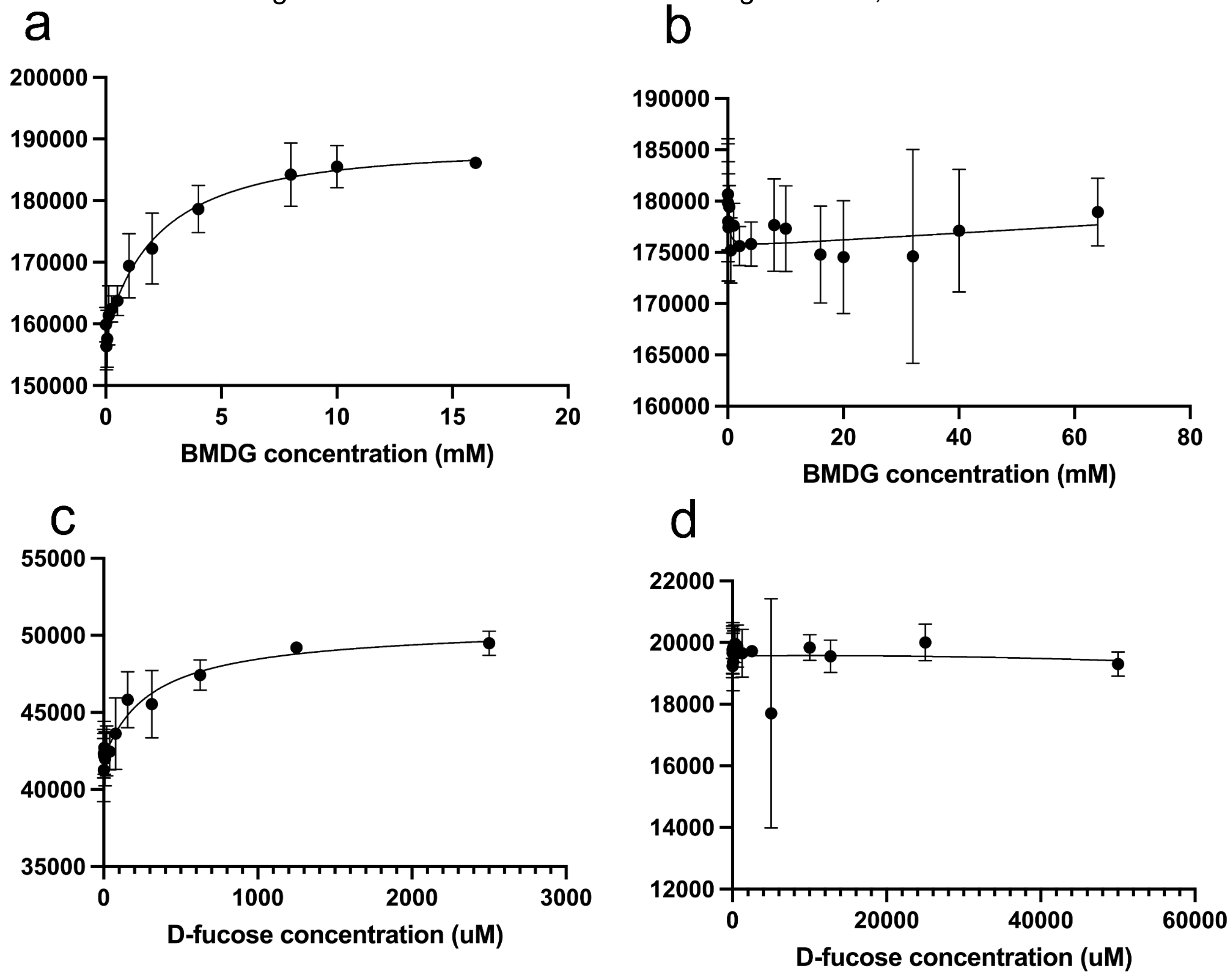

**Supplementary Figure 19.** LigPlot diagram of Anc4 and BMDG, showing hydrogen bonds and hydrophobic interactions.

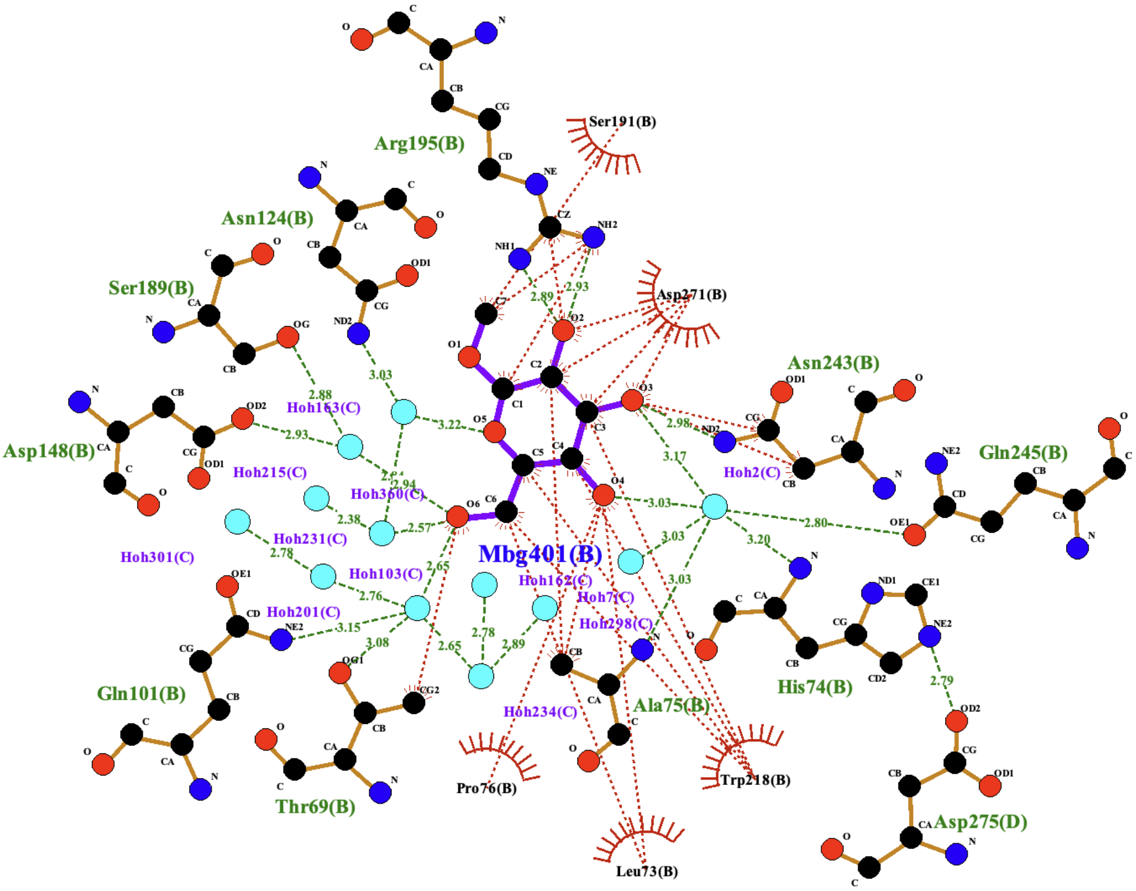

**Supplementary Figure 20.** Overlay of ΔAnc1:ᴅ-fucose (orange) and ΔAnc4:BMDG (teal).

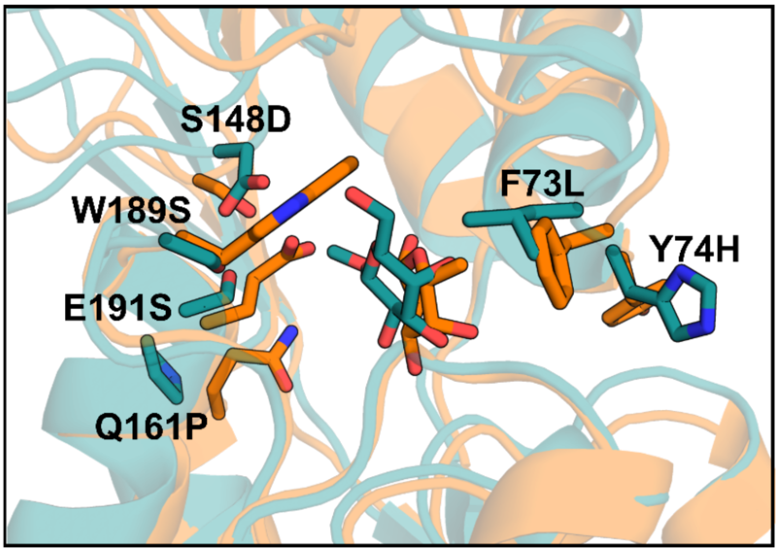

**Supplementary Table 4.** Average B-factors (Å2) for each molecule of D-fucose in Anc1:D-fucose complex.

| Chain ID | Average B-factor (Å$^2$) |
|---|---|
| A | 47.71 |
| B | 48.95 |
| C | 45.75 |
| D | 41.38 |
| E | 54.06 |
| G | 56.88 |
| H | 49.30 |
| W | 53.38 |

**Supplementary Figure 21.** Binding pockets of ancestral LGF members with omit electron density ($F_o$–$F_c$) shown in blue mesh contoured to 2σ, waters within 4 Å shown as red spheres. **a.** ΔAnc1 bound to glycerol **b.** ΔAnc1 bound to D-fucose **c.** ΔAnc4 bound to BMDG.

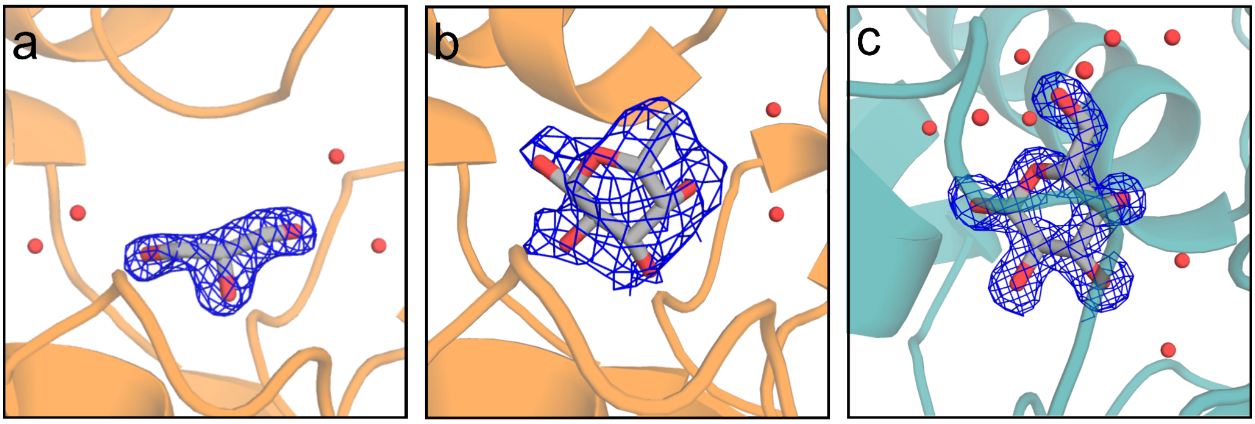

**Supplementary Table 5.** Ensemble refinement statistics

| **phenix.ensemble_refinement** | Anc1 apo | Anc1:D-fucose | Anc1:glycerol | Anc4:BMDG |
|---|---|---|---|---|
| **$\rho_{TLS}$ (%)** | 1 (optimized) | 1 (optimized) | 1 (optimized) | 1 (optimized) |
| **$\tau_{BATH}$ (K)** | 300 | 300 | 300 | 300 |
| **$\tau_{X}$ (ps)** | 0.3 | 0.4 | 0.7 | 0.9 |
| **No. of models** | 20 | 20 | 29 | 50 |
| **$R_{work}$** | 0.2114 | 0.2305 | 0.2301 | 0.1722 |
| **$R_{free}$** | 0.2874 | 0.2957 | 0.2621 | 0.2134 |
| **RMS (bonds) (Å)** | 0.017 | 0.017 | 0.017 | 0.01685 |
| **RMS (angles) (°)** | 1.41 | 1.65 | 1.47 | 1.6131 |
